# Trabecular structure correlates with leaping distance in tamarins

**DOI:** 10.1101/2025.08.30.673112

**Authors:** Uyen Nguyen, Fabio Alfieri, Alessio Veneziano, Annika Licht, John A. Nyakatura

**Author notes:** These authors contributed equally (Uyen Nguyen, Fabio Alfieri).

## Abstract

**Objectives:** The intricate trabecular architecture of long-bone epiphyses underpins functional adaptations for diverse mammalian locomotion. Despite extensive study in other mammals, tamarin trabecular structure and fine-grained differences among leaping taxa remain poorly characterized.

**Materials and Methods:** We examined humeral and tibial trabecular networks in four tamarin species representing short- and long-distance leapers using µCT-scans and a whole-epiphysis approach. We quantified network complexity with topological indices (node density, tortuosity, trabecular length, fractal dimension) alongside traditional metrics (degree of anisotropy [DA], bone volume fraction [BV/TV]) to capture both geometric and topological features.

**Results:** Long leapers exhibit significantly higher node density in both humeral and tibial epiphyses and increased trabecular tortuosity in the distal humerus. Elevated node density localizes beneath the humero-scapular joint, within the proximal humerus, and in variable regions across other epiphyses. Other parameters (DA, BV/TV, trabecular length, fractal dimension) showed no leaping-related differences, instead correlating with sex and captivity.

**Discussion:** Increased mechanical strain during longer leaps likely drives higher node density and, to a lesser extent, tortuosity in humeral and tibial epiphyses, with node density showing the strongest functional signal. While sex and captivity influence other trabecular traits, patterns in these key metrics support locomotor adaptation. Integrating whole-epiphysis analyses with novel topological indices enhances detection of subtle functional signals and complements VOI-based and traditional frameworks in comparative trabecular bone studies.

## 1. INTRODUCTION

As shown both experimentally and computationally, trabecular bone is able to respond to mechanical stimuli by osteoblast_-_driven deposition and osteoclast_-_driven resorption (Biewener et al., 1996; Keaveny et al., 2001; Kivell, 2016 and references), i.e, through modelling (*sensu* Barak, 2019). Compared to external bone shape, trabecular structure may adapt more readily to ecological and locomotor demands, both ontogenetically and evolutionarily (Kivell, 2016 and references; Alfieri et al., 2023). Numerous studies link trabecular metrics to locomotor habits (Amson et al., 2017; Arias_-_Martorell et al., 2021; Georgiou et al., 2019; Mielke et al., 2018; Ryan & Shaw, 2012; Saparin et al., 2011), yet others do not (Alfieri et al., 2025; Carlson et al., 2008), possibly due to genetic, mass, phylogenetic, hormonal, or ontogenetic influences (Havill et al., 2010; Paternoster et al., 2013; Turner et al., 2000; Doube et al., 2011; Kivell, 2016 and references; Tsegai et al., 2013; Parkinson & Fazzalari, 2012; Sowińska_-_Przepiera et al., 2024; Saers, 2017; Tsegai et al., 2018).

Methodological factors also drive conflicting results. Trabecular isolation via regularly shaped Volumes of Interest (VOIs) offers reproducibility and lower computation but biases may derive from VOI size, shape, and location (Fajardo & Müller, 2001; Ryan & Ketcham, 2001; Kivell et al., 2011; Lazenby et al., 2011). It led to develop whole_-_epiphysis methods (Bachmann et al., 2022; Veneziano et al., 2021). Parameter choice represents another potential bias: e.g. Degree of Anisotropy (DA) and Bone Volume Fraction (BV/TV) (‘traditional parameters’ hereafter) are deemed most informative (Maquer et al., 2015; Stauber et al., 2006) and often used exclusively (Arias_-_Martorell et al., 2021; Ryan & Ketcham, 2001), though topological indices (i.e. measured on topological skeletons, that only preserve trabecular node–branch relationships) have emerged (Alfieri et al., 2025; Veneziano et al., 2021). Moreover, results depend on the studied skeletal region—e.g. femoral trabeculae may be more locomotor_-_informative than humeral (Ryan & Shaw, 2012)— and sample, e.g. positive results often yielded by studying hominids (Arias_-_Martorell et al., 2021; Georgiou et al., 2019; Kivell et al., 2018; Tsegai et al., 2013), contrasting studies of broader phylogenetic contexts (Gonet et al., 2022; Ryan & Shaw, 2012; Ryan & Walker, 2010).

The study of leaping primates has dealt with these aspects. Leaping strepsirrhines yield distinctive femoral head DA and BV/TV in cubic VOIs, a model later applied to omomyids and tarsiers too using spherical VOIs (Ryan & Ketcham, 2001; 2002; 2005). Although finite element models did not confirm the biomechanical optimization of leaping primate trabecular bone (Ryan & von Rietbergen, 2005), leaping—evolved multiple times (strepsirrhines, tarsiers, platyrrhines), imposing strong limb loadings (Demes et al., 1999), and pivotal for primate evolutionary inferences (Boyer et al., 2017)—remains an instructive eco_-_morphological system.

Comparative trabecular analyses of leaping platyrrhines appeared only recently when Berles et al. (2024) measured VOI_-_based metrics in four tamarin species (*Saguinus midas*, *S. mystax, S. imperator*, *Leontocebus nigrifrons*), all using arboreal quadrupedalism besides leaping (Garber, 1991; Garber & Pruetz, 1995; Nyakatura & Heymann, 2010). Despite ecological and locomotor differences among them, long bone morphology of the four taxa did not segregate them accordingly (Berles et al. 2024). However, it also arose that *Leontocebus nigrifrons* and *Saguinus mystax* leap farther than *S. imperator* and *S. midas* (hence termed ‘long leapers’ and ‘short leapers’, respectively) (Berles et al., 2024) and long leapers showed slightly higher BV/TV and DA, suggesting trabecular sensitivity to leaping distance (Berles et al., 2024). Most eco_-_morphological trabecular studies have contrast distinct locomotor categories (e.g., Amson & Kilbourne, 2019; Assif & Chirchir, 2024; Ingle & Porter, 2021; Ryan & Shaw, 2012; Saparin et al., 2011), while few probe subtle variation (and typically in apes, Harper & Patel, 2024; Deckers et al., 2022; Ragni, 2020), questioning trabecular bone’s resolution of fine locomotor differences.

Assuming that Berles et al.’s weak locomotor trend in trabecular bone reflects a stronger signal attenuated by VOI approach and traditional metrics, we reanalyzed the same individuals using a whole_-_epiphysis approach (Veneziano et al., 2021), though both traditional and topological traits. The biomechanical meaning of some traditional parameters is well established. DA quantifies trabecular alignment; directionally stereotyped locomotion expectedly produces higher anisotropy (Harrigan & Mann, 1984; Ryan & Ketcham, 2001). BV/TV measures the inner epiphyseal trabecular bone proportion and is hypothesized to increase in response to higher loadings (Arias-Martorell et al., 2021). Topological indices lack direct biomechanical validation but their geometric correlates were related to mechanical loadings. Node density (NodDen) parallels BV/TV and correlates with compressive strength, so increased loading should raise NodDen (Cendre et al., 1999; Veneziano et al., 2021). Trabecular length (TrabLen) should inversely relate to strength, since denser trabeculae (i.e, higher NodDen) are also expected to be shorter (Parkinson et al., 2012). Trabecular tortuosity (TrabTort, the arc/Euclidean length ratio) governs elasticity: higher TrabTort is hypothesized to yield more elastic yet load-resistant structures (Fyhrie & Zauel, 2015; Roque et al., 2012; Roque & Alberich-Bayarri, 2015). Fractal dimension (FD) reflects self-similarity across scales. High FD with low density being associated with mechanical inactivity (Feder, 1988; Pornprasertsuk et al., 2001; Ammann & Rizzoli, 2003; Feltrin et al., 2004; Messent et al., 2005a,b), FD should inversely correlate with loading (Note S1, Supplementary Material; Alfieri et al., 2025; Veneziano et al., 2021).

We re-examined *Leontocebus nigrifrons*, *Saguinus mystax*, *Saguinus imperator*, and *Saguinus midas* through a whole epiphysis approach and additionally quantifying topological traits. Since their leaping distance categories do not align with phylogeny and they share similar body masses (≈ 400–600 g; Garber & Teaford, 1986), we assumed that allometric and phylogenetic effects were minimized. We sampled humeral and tibial epiphyses because hindlimbs generate propulsion in take-off (Channon et al., 2010; Demes et al., 1995; 1999), and forelimbs absorb landing forces (Berles et al., 2022; Garber, 1991; Garber et al., 2012). The tibia, as the primary lower-leg load bearer, directly transmits ground reaction forces and is pivotal for shock absorption (Bourne et al., 2018).

Small primates, especially generalized, leapers, incur exceptionally high leaping forces (Demes et al., 1999). Also, longer leaps generate greater propulsive and landing forces (Demes et al., 1999; Nauwelaerts & Aerts, 2006). Tamarins being small generalized leapers and consistently with functional predictions for trabecular variables, we predict higher BV/TV, NodDen, and TrabTort, and lower TrabLen and FD in humeral and tibial epiphyses of ‘long leapers’, compared to ‘short leapers’. While no directional differences are involved in long vs. short leaping, stronger loads in long leapers may still result in more anisotropic trabeculae, i.e. higher DA.

## 2. MATERIALS AND METHODS

### 2.1 Micro-CT Scanning

We studied twelve humeri and tibiae, from the same individuals studied in Berles et al. (2024). They represent *Leontocebus nigrifrons, Saguinus midas, S. imperator,* and *S. mystax*, were sourced from the American Museum of Natural History (AMNH, New York) and the Field Museum of Natural History (FMNH, Chicago, IL) and imaged via µCT on a Nikon Metrology XT H 225 ST (voxel=0.0165–0.018 mm, 100 kV, 165 µA, 0.2 mm Cu filter) at Duke SMiF, and on a GE phoenix vtomex s (voxel=0.0154 mm, 170–172 kV, 79–85 µA, no filter) at UChicago PaleoCT (Table S1). Scans are downloadable from MorphoSource. Although three individuals were likely subadults, we only analyzed fully fused epiphyses (Table 1).

**Table 1:**
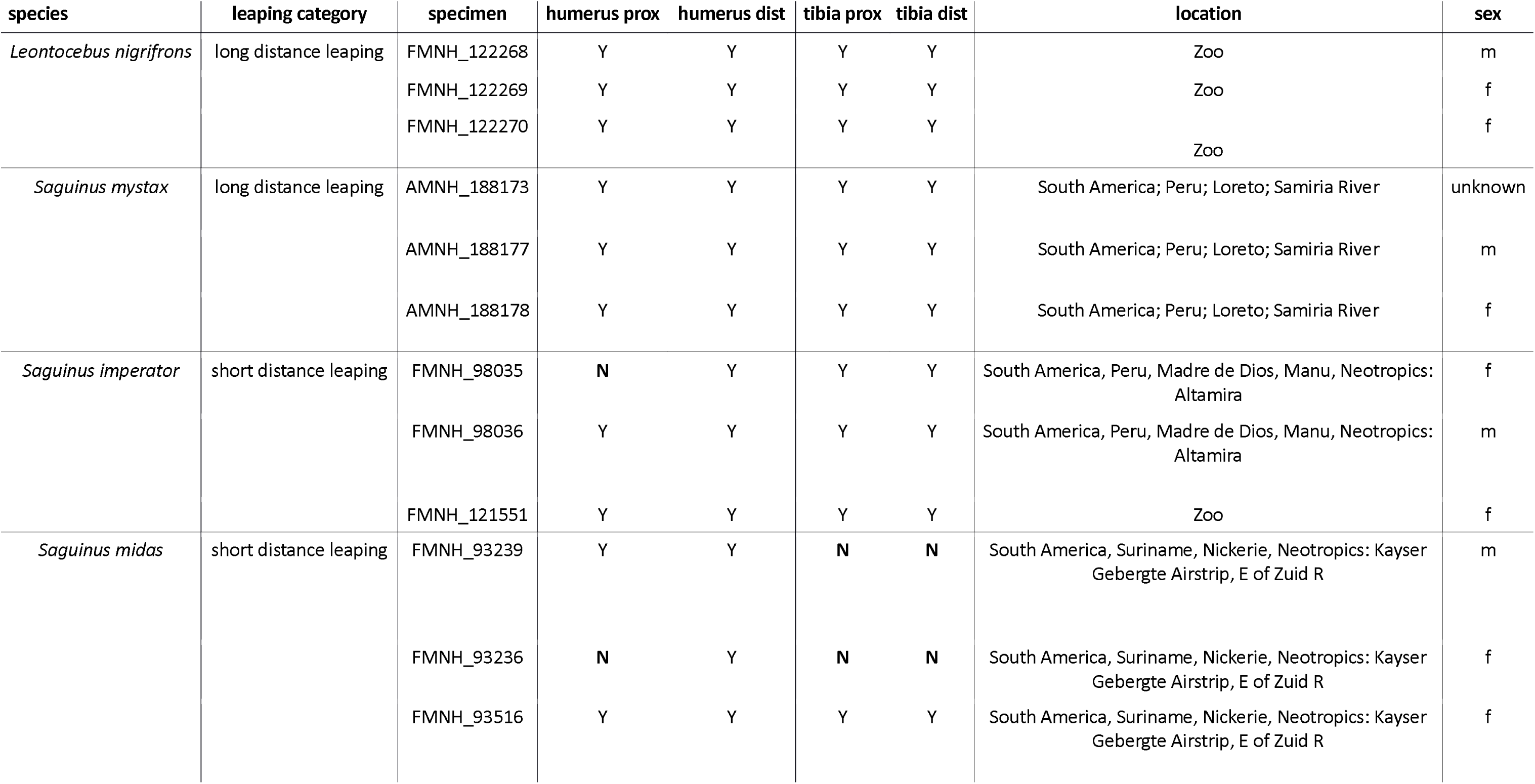
Specimens studied in this work, together with information on the species to which they belong, their leaping category (i.e., leaping distance), and epiphyseal elements that were included (as indicated with Y) or discarded (as indicated with N) in case of not fully fused epiphysis. Prox and dist are abbreviations for proximal and distal, respectively, information on the location of collection and sex.

Three individuals per species limit intraspecific variation assessments. Yet, we calculated coefficients of variation (CV; Table S4). Potential effects of captivity and sex differences, both involved in our sample (Table 1), were visualized through Principal Component Analysis (Notes S4–S5).

### 2.2 Trabecular bone isolation

Using VG Studio Max 3.3 (Volume Graphics, Heidelberg, Germany), we aligned all the humeri and tibiae in the same standard position, through a protocol mostly following Ruff (2002) and detailed in Notes S1-S2. On oriented bones, by using the same anatomical markers across specimens, the epiphyses were cropped. For the proximal humerus, we used the most proximal point of the head and the most distal level of the anatomical neck. For the distal humerus, we used the most proximal point of the capitulum and the most distal point of the bone. For the proximal tibia, we took the most proximal point of the bone and the most distal point of the lateral condyle. For the distal tibia, we took the first point where the medial malleolus can be detected (going proximodistally along the image stack) and the most distal point of the bone (Fig. 2). To isolate trabecular bone, we followed Veneziano et al. (2021). In brief, image stacks were binarized in ImageJ (Schneider et al., 2012) (‘Threshold’ tool), hence separating bone and non-bone voxels. Thresholds were manually adjusted after visual comparison with original scans to avoid misclassification. For binarized epiphyses, we automatically isolated trabecular bone exploiting the ‘indianaBones’ R (v4.3.2; R Core Team, 2023) package (https://github.com/AlessioVeneziano/IndianaBones), obtaining trabecular bone from whole epiphyses (Region of Interest, ROI). The isolation algorithm was iterated until visual comparison to stacks including cortical bone too, confirmed optimal isolation (Fig. 2).

**Figure 1:**
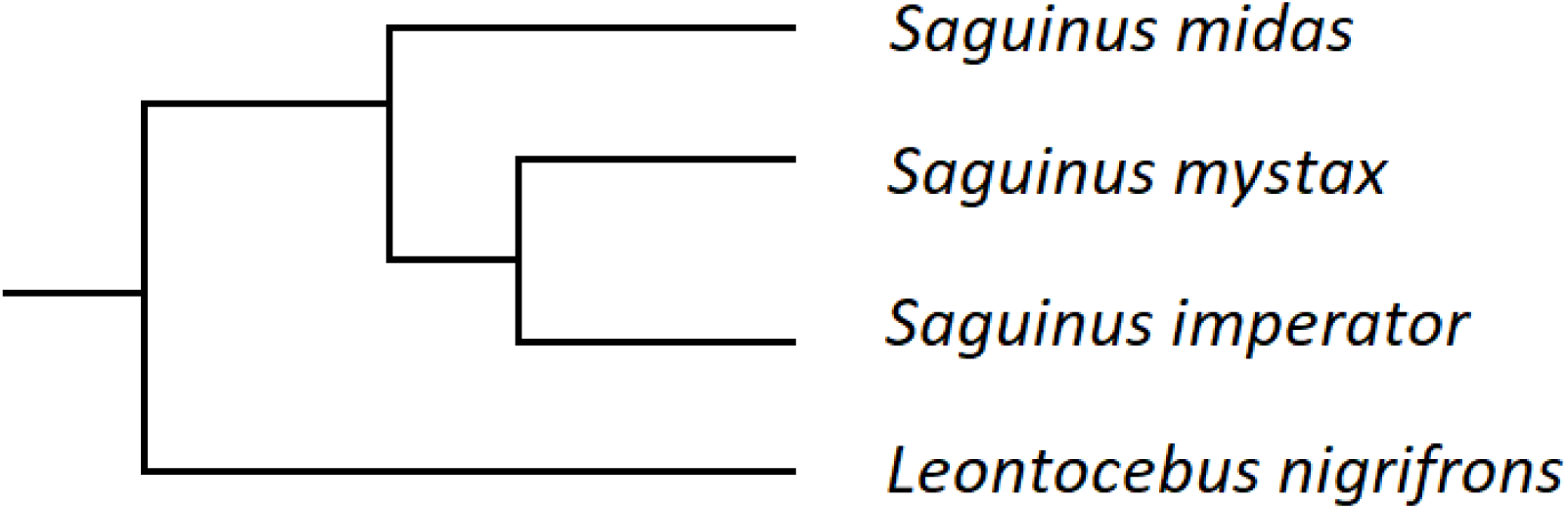
Topology of the phylogenetic relationships among the four examined tamarin species

**Figure 2:**
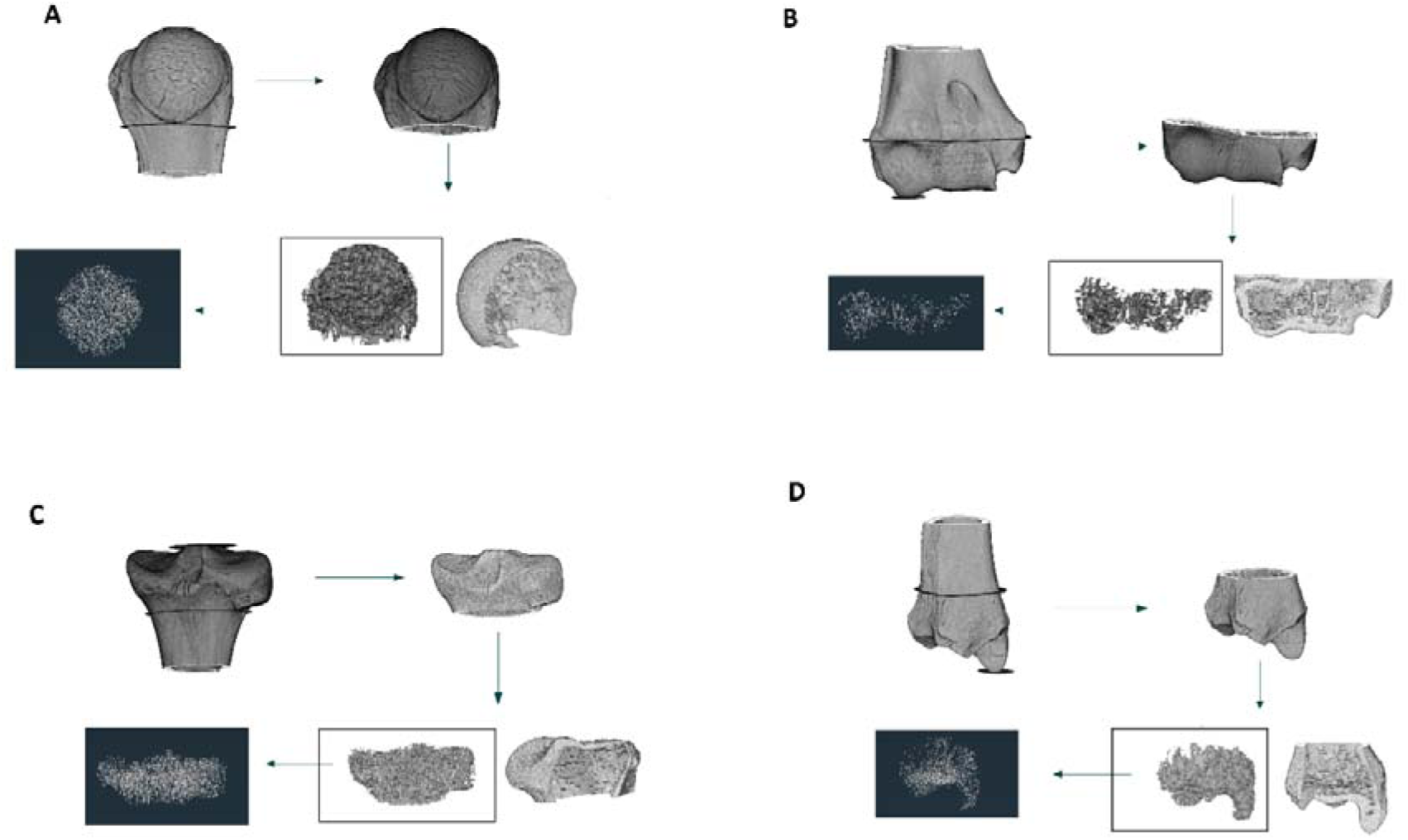
Trabecular bone isolation is summarized for the humeral (A. proximal; B. distal) and the tibial (C. proximal; D. distal) whole epiphyses of S. mystax AMNH 188173. From fixed anatomical markers on proximal (first step in A, C) and distal epiphyses (first step in B, D), we cropped the epiphyses (second step in A-D), we isolated the ROIs (through the exclusion of cortical bone; third step in A-D) and we transformed them into topological skeletons (four step in A-D).

### 2.3 Quantification of trabecular traits

ROIs were skeletonized in Amira 6.0.0 (Auto Skeleton tool), reducing the trabecular network to topological skeletons. In R we extract topological indices from topological skeletons (see Alfieri et al. 2025; R code): NodDen (mean and median), TrabTort (mean and median), TrabLen (mean) and FD. Using both mean and median for NodDen and TrabTort is intended to capture outlier-driven variability at volume margins. We also generated NodDen heatmaps (Notes S6–S9) to visualize spatial distribution across the epiphysis. For details on topological indices, see Note S1, Veneziano et al. 2021 (fig. 3) and Alfieri et al. 2025. The traditional parameters BV/TV and DA were measured on ROIs using BoneJ (Doube et al., 2010, ImageJ plugin). Since the TV computed by BoneJ is the full image_-_stack volume rather than the ROI volume, we used the ROI volume from the ‘indianaBones’ isolation as TV, and divided BoneJ’s bone_-_voxel count (BV) by it to calculate BV/TV.

**Figure 3:**
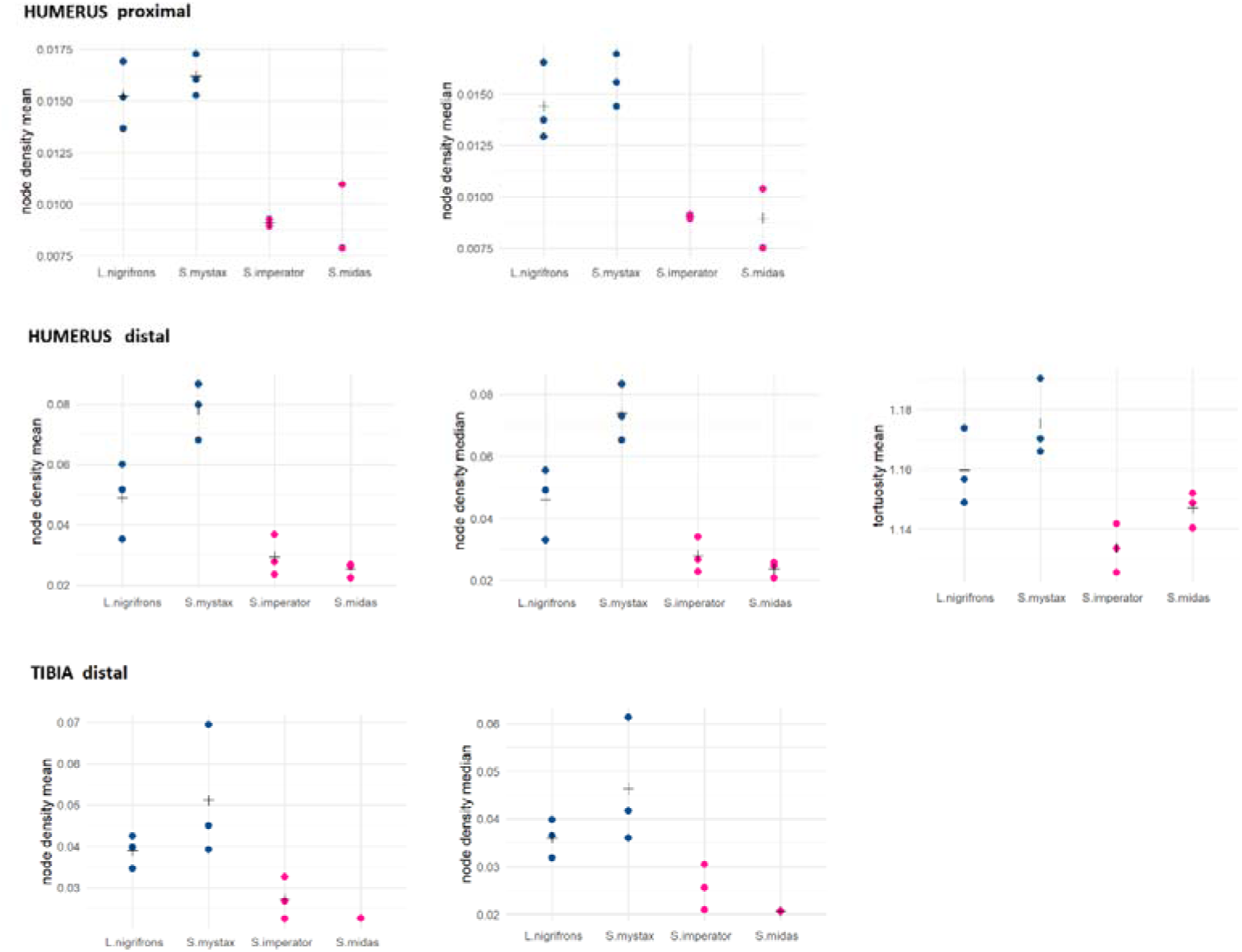
Scatterplots of the topological indices that significantly discriminate long leapers (blue observations) and short leapers (purple observations). + = mean values.

### 2.4 Statistical analysis

Because the four closely related taxa do not cluster by leaping distance (Fig. 1; Table 1), we assumed negligible phylogenetic effects and did not apply phylogenetically informed tests.

Also similar body mass data led us to assume negligible allometric effects. Comparable humerus (mean 52.94 ± 3.19 mm) and tibia (mean 68.87 ± 5.27 mm) lengths (Table S1) further support size homogeneity.

Due to low sample size, we used nonparametric Mann–Whitney U tests (α = 0.5) to compare each trabecular variable between long and short leapers (Table 2). P-values were corrected by False Discovery Rate (FDR) to account to control the expected proportion of false positives deriving from multiple testing. FDR was preferred over other more conservative methods (e.g., Bonferroni) due to the preliminary nature of the study. Differences between long and short leaping taxa were visualized in scatterplots (Figs. 3–5).

**Figure 4:**
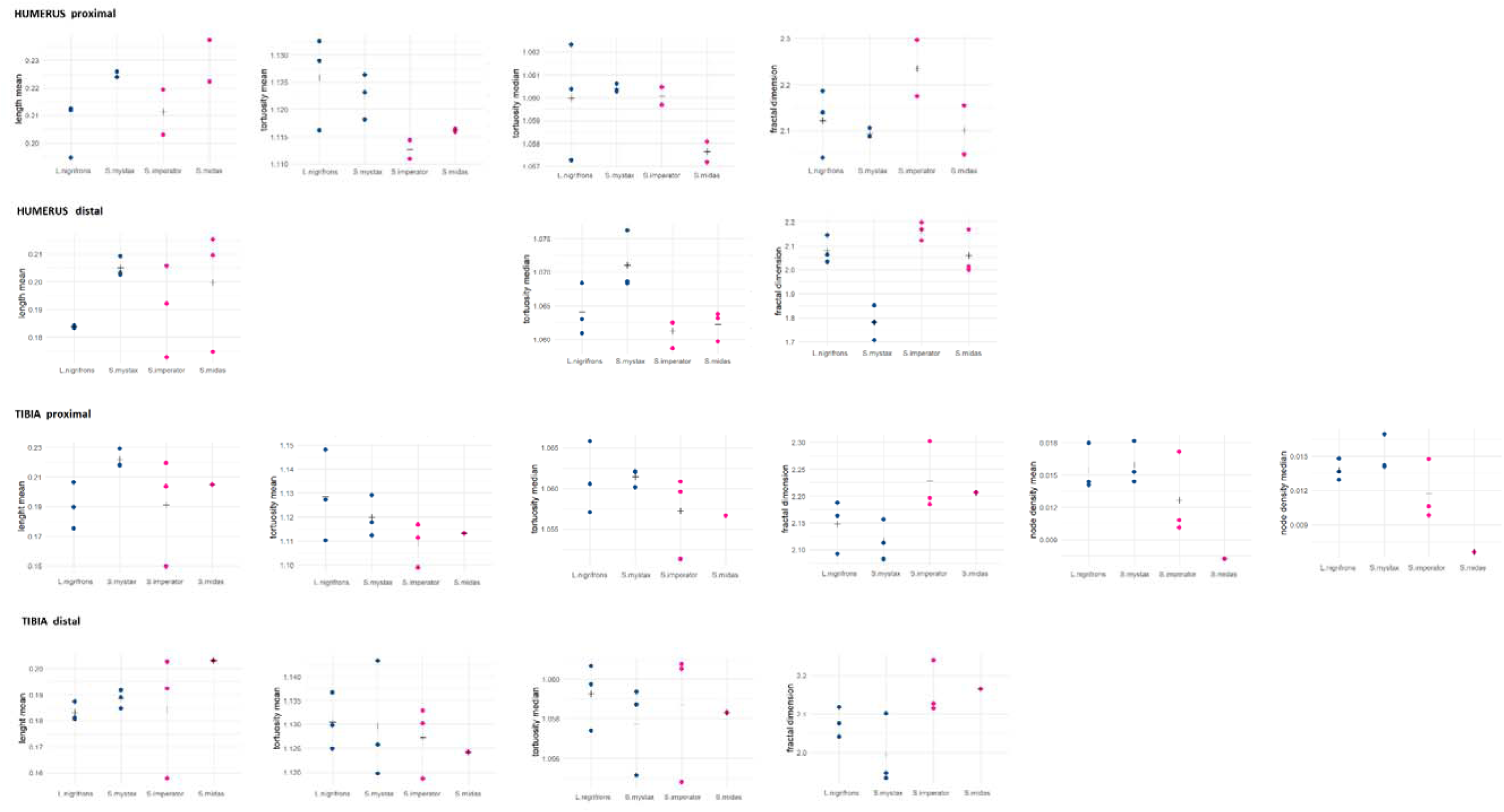
Scatterplots of the topological indices that did not discriminate long leapers (blue observations) and short 869 leapers (purple observations) values.

**Figure 5:**
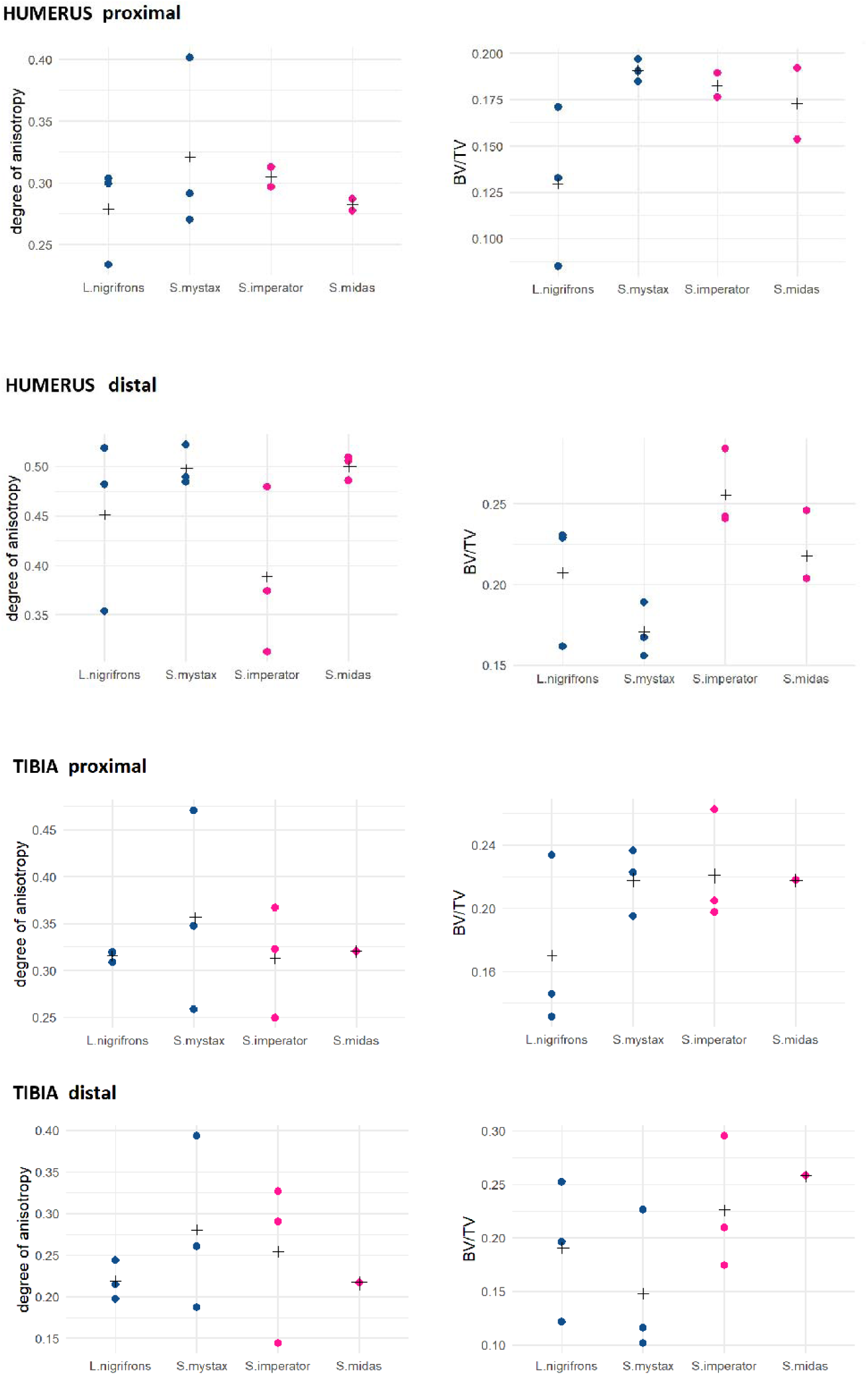
Scatterplots of the traditional parameters for long leapers (blue observations) and short leapers (purple observations). + = mean values.

**Table 2:**
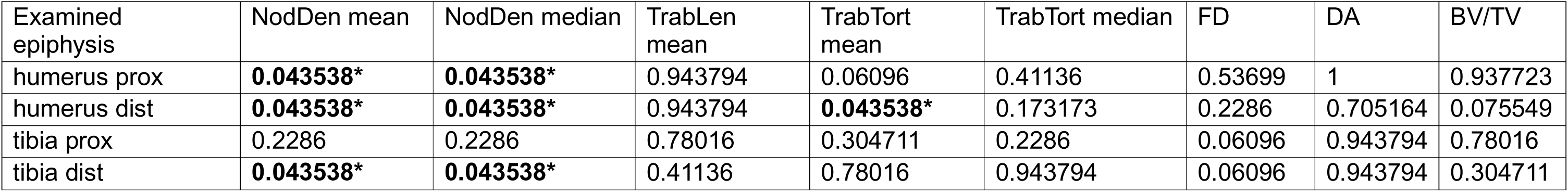
For each anatomical region studied here, i.e., examined epiphyses, (in rows), and for each topological index of complexity there computed (in columns), the p-value resulting from testing for a significant difference between long distance leaping and short distance leaping tamarins is shown. See Methods and above for details and abbreviations. The values were obtained with the Mann-Whitney-U-Test/Wilcoxon-Rank-Sum-Test and after fdr correction.

## 3. RESULTS

Significant correlations between trabecular traits and leaping distance appear across both bones and nearly all epiphyses. In three of four epiphyses (excluding proximal tibia), long leapers show significantly denser trabeculae (higher NodDen, mean and median) that, in the distal humerus, are also more tortuous (higher mean TrabTort). No other parameters differed following leaping distance (Table 2; Figs. 3–5).

Concentrated higher NodDen regions are present beneath the humeral head articulation with glenoid cavity and variably in distal humerus and tibia for long leapers, whereas short leapers display more uniform NodDen distributions (Notes S6–S9)

PCAs suggest that females exhibit more trabeculae that are more complex (higher FD) in both bones, shorter (lower TrabLen) in the humerus, more curved (higher TrabTort), less preferentially aligned (DA) and with more bone volume fraction (BV/TV) in the tibia. Captive individuals show less trabeculae that are tortuous (lower TrabLen) and more complex (higher FD) in the proximal humerus, with higher TrabTort in the proximal tibia, and lower BV/TV in both proximal epiphyses. Noticeably, these patterns—i.e., which trait from which epiphysis— mostly differ from those associated with leaping distance, with the only exception of distal tibia NodDen (Notes S4-S5).

As for intraspecific variation, CV overall ranges 0.00017–0.46 (mean 0.096), while for only for leaping distance_-_related traits it ranges 0.005–0.31 (mean 0.134).

## 4. DISCUSSION

Leaping demands anatomical adaptations enabling to generate high-power over short intervals (Mo et al. 2020). In primates, both specialist (galagids, tarsiers) and generalized leapers (lemurs, platyrrhines) exhibit these adaptations, e.g. robust, elongated femora, with enlarged heads, filled with anisotropic trabecular architecture (Connour et al. 2000; Polvadore et al. 2024; Ryan & Ketcham 2001). Despite kinetic data on differential force production among leaping taxa are present (Demes et al. 1996; 1999), few studies have compared morphological variation between differently leaping primates.

Recently, Berles et al. (2024) studied leaping tamarins—*L. nigrifrons, S. mystax, S. imperator, S. midas*—that differ in leaping distance (Garber 1991; Garber & Pruetz 1995; Nyakatura & Heymann 2010). Although not significantly, long leapers (*L. nigrifrons, S. mystax*) showed modestly higher DA and BV/TV than short leapers (*S. imperator, S. midas*), possibly reflecting greater take-off and landing forces in primates leaping over longer distances (Demes et al. 1999).

We hypothesized that in Berles et al. 2024, VOI_-_based sampling and exclusive reliance on traditional metrics attenuated stronger signals. Hence, by re_-_analysing the same individuals, we studied whole humeral and tibial epiphyses and quantified both traditional and topological variables, aiming to capture fine_-_scale trabecular variability. As detailed below, we identified trabecular correlates to leaping distance, while other aspects were identified as affected by non-functional factors, i.e. sex, and captivity.

### 4.1 Covariation of trabecular node density and tortuosity with leaping distance

NodDen appears to be the trabecular feature with the strongest signal related to leaping distance. Indeed, it extensively - i.e. in proximal/distal humerus and the in distal tibia - yielded significant differences between long, i.e. *L. nigrifrons* and *S. mystax*, and short leapers, i.e. *S. midas* and *S. imperator* (though both mean and the median, Table 2, Fig. 3). Hence, in long distance leapers the most common way through which the forelimb and hindlimb trabecular structure possibly adapts to higher loads deriving from longer leaps (Demes et al. 1999) appears to be an increase in node density. This index directly derives from density measures and, in turn, represents a concept parallel to trabecular bone compactness, i.e. BV/TV (see e.g. Tsegai et al. 2018 and Note S1). Higher loadings on joints are expected to increase bone inner compactness and density (e.g. Arias-Martorell et al. 2021 for BV/TV, see also similar density-related measures positively related to maximum compressive strength, Cendre et al. 1999). Higher NodDen in long leapers, due to expected higher loadings associated with longer leaping distances (Demes et al. 1999), supports this hypothesis. Also, the fact that this pattern arose in both the bones and for both proximal and distal epiphyses, is in agreement with our expectation of generalized increased loadings in all the studied epiphyses.

Long leapers tend to have isolated areas of high NodDen in the epiphyses, that likely cause the significantly higher values and can be anatomically defined especially in the humeral head, i.e. surface articulating with the scapula. This condition is less common in short leapers, conversely showing more spatially uniform NodDen values (Note S6-S9). The humeral trabecular bone underlying the glenohumeral joint seems to be the region more consistently showing an increase in NodDen in long leapers (Note S6). The primary role covered by glenohumeral joint in transmitting and dissipating loadings in primate forelimb (Arias-Martorell et al. 2015, Arias-Martorell, 2019, Preuschoft et al. 2010), is in agreement with the mechanical meaning of higher NodDen that we propose for long leapers’ humeri. As for the other epiphyses, the regions of higher NodDen are less regularly concentrated in the same sub-regions and are hence more challenging to be interpreted. Pending more detailed kinematics data on the studied species that would allow to expect higher density in specific portions of the epiphyses, we hypothesize that the higher loadings deriving from longer leaping distances are transmitted throughout the epiphyses, resulting in high NodDen zones spread within the trabecular network.

Another parameter that yielded significant correlation with leaping distance is TrabTort, that is significantly higher in long leapers’ distal humerus (Table S2). Similarly to NodDen (see above), this pattern is in agreement with our functional expectations. Indeed, the extent to which trabeculae are curved inversely relates to stiffness (Roque et al. 2012; Roque and Alberich-Bayarri 2015), i.e. more tortuous trabeculae correspond to structurally more elastic networks, which potentially represents an adaptation to higher loadings. However, we interpret the trend found for TrabTort more cautiously compared to what we discussed above for NodDen. First, this result only derives from a single epiphysis, i.e. distal humerus, and only with mean values. Also, the absence of heat maps as informative as those for NodDen makes interpretation more challenging at this stage, i.e. we cannot determine whether the distal humeral trabeculae of long leapers are more convoluted throughout the entire epiphysis or only in specific sub-regions.

### 4.2 Other factors affecting tamarin trabecular structure

Besides the patterns discussed above for NodDen and TrabTort, we did not find any other significant relationship with leaping distance for the other variables (Table 2, Figs. 4-5). While the sample composition of this work allowed us to reasonably exclude allometric and phylogenetic effects on the studied traits (see Methods), we identified some potential effects of sex and captivity on trabecular variables (Notes S4-S5). This is not surprising, since effects of sex and captivity on trabecular structure have been previously reported (e.g. Eckstein et al. 2007, Zack et al. 2022). Remarkably, all the potentially sex and captivity related patterns that we highlighted do not concern the variables for which we hypothesized a mechanical meaning. The only exceptions are possible effects of sex in distal tibia NodDen (Note S4). However, the contribution of NodDen to the discrimination of male from female individuals appears to be due to extremely high NodDen values of one single specimen, i.e. *Saguinus mystax* AMNH 188177 (see NodDen vectors direction in distal tibia variable loading plots and PCA biplots, Note S4). This individual is not a sub-adult, is wild caught (hence potential age and captivity-related biases can be excluded). Yet, it belongs to a long-distance leaping species (Table 2) and, thus, its extremely high NodDen would be consistent with higher loadings (see above). A potential effect of sex and captivity was yielded by TrabTort too. Although it occurs in the tibia (and we found a relationship with leaping distance in the distal humerus) it is another element which led us to temper our functional interpretation of TrabTort (see above).

A striking result is the absence of correlation between DA and BV/TV with leaping distance. Indeed, the two minor trends of positive covariation between these two variables and leaping distance found by Berles et al. stimulated our work. As for DA, a possible explanation lies in the fact that this parameter measures a local property, i.e. the degree of preferential alignment (Harrigan and Mann, 1984). Hence, over whole epiphyses, the signal of DA may be less clear. As for BV/TV absence of relationship with leaping distance, the pattern is more puzzling to be interpreted, especially considering that it is conceptually related to NodDen (i.e. the parameter yielding the strongest signal related to leaping distance, see above). It should be noted that, although the two parameters measure similar properties, they are also computationally different (i.e. NodDen is computed on topological skeletons). This discrepancy may emphasize the need for combined utilization of both these parameters in comparative analyses of trabecular bone.

Due to the relatively novel methodological approach employed in this work and the scarcity of previous analyses of tamarin trabecular structure, there is not a broad comparative framework that we can exploit to interpret our outcomes (except for Berles et al. 2024, discussed above). The same techniques were recently employed in an analysis of primate distal fibula trabecular structure (Alfieri et al. 2025). Although no main co-variation with locomotion was found, one of the parameters here found to potentially relate to leaping distance, i.e. TrabTort, was proposed to show a relationship with human bipedalism (Alfieri et al. 2025)

The small sample size represents a limitation of this work. Indeed, it only allowed us to preliminarily highlight patterns that further dedicated studies need to confirm. One of the main issues caused by our sample size is the low representation of individuals within a species, which limits our possibility to estimate intraspecific variation. For instance, tibial patterns for *S. midas* should be interpreted with caution, since the species is represented by a single specimen (due to exclusion of tibiae with unfused epiphyses). We were however able to estimate intraspecific variability by computing CV for each species and trait: in general, a CV reaching 0.46 suggests a substantial intraspecific variability (although the average value indicates an overall moderate effect, i.e. 0.096). Noticeably, if only the traits significantly related to leaping distance are considered, the maximum CV decreases to 0.31, with an average value again suggesting moderate intraspecific variability for the functional relationships that we discuss. Another potential bias may derive from the automatic trabecular bone isolation procedure that may involve unwanted inclusion of cortical bone in the studied network. It, in turn, may affect some variables (e.g. node density can be artificially increased if cortical bone is mistakenly included). However, we carefully assessed the results of each automatic trabecular isolation, checking for the potential presence of cortical bone, and we can reasonably exclude that substantial regions of cortical bone were included in the studied ROIs (see also an example of excluded cortical fraction in Fig. 2).

### 4.3 Advantage of the whole-epiphysis approach and computation of topological indices

Our outcomes suggest that employing a whole epiphysis approach and computing topological indices, may allow to identify patterns potentially reflecting fine-grained locomotor differences, i.e. different leaping distances. Berles et al., examining the same specimens, could not find significant differences in BV/TV (functionally comparable to node density, see above) and DA between short and long-distance leapers (Berles et al., 2024). While it suggests that a whole epiphysis approach may be preferred to increase the detection of locomotor signal in trabecular bone, one should also note that several advantages are still involved in VOI approaches. Besides the easier reproducibility and low computation demand, VOIs allow to better represent regional properties (e.g. DA, see discussion above) (Kivell et al. 2011). It can explain why the minor trend of co-variation between DA and leaping distance in Berles et al. 2024 was neither found as significant nor detected qualitatively in this work.

## 5 CONCLUSIONS

By employing a whole-epiphysis approach, and computing both traditional trabecular parameters and topological indices, we characterized the relationship between humeral/tibial trabecular structure and leaping distance of tamarins. Our aim was to obtain information on possible adaptations of the trabecular structure related to minor differences in leaping style. Results for two traits, i.e. node density, especially, and trabecular tortuosity, appears to be consistent with our expectations: long distance leapers, i.e*., L. nigrifrons* and *S. mystax,* show higher trabecular node density (NodDen) and trabecular tortuosity (TrabTort) compared to short distance leapers, i.e., *S. midas* and *S. imperator*. These patterns possibly relate to higher loadings expected in long bone epiphyses of species leaping across longer distances. We propose that these two specific properties of tamarin humeral/tibial trabecular bone are associated with leaping distance. As for the other trabecular properties that we computed, they did not yield any relationship with locomotor aspects and were likely associated with non-functional factors, as sex and captivity. We suggest that a whole-epiphysis approach, combined with the computation of topological indices, offers a powerful framework for characterizing trabecular structure. However, traditional parameters and VOI-based analyses may still reveal additional, complementary patterns and should not be overlooked.

## Author contributions

FA and JAN conceived of the study; FA processed CT-scan data; UN and AL extracted and analyzed raw data; AV supervised trabecular bone virtual isolation and computation of topological indices. UN and FA drafted the manuscript and figures. All authors interpreted the results and commented on the initial draft.

## Acknowledgements

We thank Léo Botton-Divet for CT data access, the Nyakatura Lab at Humboldt-Universität zu Berlin, and Eli Amson for methodological support and insightful discussion. This work used the Duke University Shared Materials Instrumentation Facility (SMIF), part of the North Carolina Research Triangle Nanotechnology Network (RTNN) supported by NSF award ECCS-2025064 under the National Nanotechnology Coordinated Infrastructure (NNCI). Funding was provided by the German Research Foundation (DFG NY 63/2-1 to JAN) and by the SNSF Postdoctoral Fellowship TMPFP3_217022 to FA.

## Conflict of interest statement

The authors have no conflicts of interest to declare.

## Data Availability Statement

Supplementary Material i.e., Notes S1 to S9 and the R code are downloadable from Figshare (https://figshare.com/s/e803d685895a41ec9f57?file=56303942) (Nguyen et al., 2025). The µCT scans of the humeri and the tibiae studied in this work are freely downloadable from MorphoSource (https://www.morphosource.org/). The R package “indianaBones” is available at https://github.com/AlessioVeneziano/IndianaBones and at https://zenodo.org/record/7615165.

## Ethical statement

No human remains are involved in this work. We studied virtual data deriving from animal remains represented by bones, which are stored and collected as dried specimens in museum collections (as detailed in the Methods section). These bones were sampled and CT-scanned for previous analyses (i.e., Berles et al., 2024) and we used these already generated virtual data to perform this analysis. Hence, our procedures to study animal remains did not require any ethical review.

## Notes

https://figshare.com/s/e803d685895a41ec9f57?file=56303942

### Competing Interest Statement

The authors have declared no competing interest.

## References

Alfieri, F. (2022). Convergent evolution of humeral and femoral functional morphology in slow arboreal mammals. PhD dissertation, Humboldt-Universität zu Berlin, Berlin, Germany. 10.18452/25436.

Alfieri, F., Botton-Divet, L., Nyakatura, J. A., & Amson, E. (2022). Integrative approach uncovers new patterns of ecomorphological convergence in slow arboreal xenarthrans. Journal of Mammalian Evolution, 29(2), 283–312. 10.1007/s10914-021-09590-5.

Alfieri, F., Botton-Divet, L., Wölfer, J., Nyakatura, J. A., & Amson, E. (2023). A macroevolutionary common-garden experiment reveals differentially evolvable bone organization levels in slow arboreal mammals. Communications Biology, 6(1), 995. 10.1038/s42003-023-05371-3.

Alfieri, F., Veneziano, A., Panetta, D., Salvadori, P. A., Amson, E., & Marchi, D. (2025). The relationship between primate distal fibula trabecular architecture and arboreality, phylogeny and size. Journal of Anatomy, 246(6), 907–935. 10.1111/joa.14195.

Ammann, P., & Rizzoli, R. (2003). Bone strength and its determinants. Osteoporosis international, 14, 13–18. 10.1007/s00198-002-1345-4.

Amson, E., & Kilbourne, B. M. (2019). Trabecular bone architecture in the stylopod epiphyses of mustelids (Mammalia, Carnivora). Royal Society Open Science, 6(10), 190938. 10.1098/rsos.190938.

Amson, E., Arnold, P., van Heteren, A. H., Canoville, A., & Nyakatura, J. A. (2017). Trabecular architecture in the forelimb epiphyses of extant xenarthrans (Mammalia). Frontiers in Zoology, 14(1), 1–17. 10.1186/s12983-017-0241-x.

Assif, L., & Chirchir, H. (2024). Trabecular bone morphology in big cats reflects the complex diversity of limb use but not home range size or daily travel distance. The Anatomical Record, 307(1), 208–222. 10.1002/ar.25302.

Arias_-_Martorell, J., Tallman, M., Potau, J. M., Bello_-_Hellegouarch, G., & Pérez_-_Pérez, A. (2015). Shape analysis of the proximal humerus in orthograde and semi_-_orthograde primates: Correlates of suspensory behavior. American Journal of Primatology, 77(1), 1–19.

Arias_-_Martorell, J. (2019). The morphology and evolutionary history of the glenohumeral joint of hominoids: A review. Ecology and Evolution, 9(1), 703–722.

Arias-Martorell, J., Zeininger, A., & Kivell, T. L. (2021). Trabecular structure of the elbow reveals divergence in knuckle-walking biomechanical strategies of African apes. Evolution, 75(11), 2959–2971.

Bachmann, S., Dunmore, C. J., Skinner, M. M., Pahr, D. H., & Synek, A. (2022). A computational framework for canonical holistic morphometric analysis of trabecular bone. Scientific Reports, 12(1), 5187.

Barak, M. M. (2019). Bone modeling or bone remodeling: That is the question. American Journal of Physical Anthropology, 172(2), 153–155.

Berles, P., Heymann, E. W., Golcher, F., & Nyakatura, J. A. (2022). Leaping and differential habitat use in sympatric tamarins in Amazonian Peru. Journal of Mammalogy, 103(1), 146–158. 10.1093/jmammal/gyab121.

Berles, P., Wölfer, J., Alfieri, F., Botton-Divet, L., Guéry, J. P., & Nyakatura, J. A. (2024). Linking morphology, performance, and habitat utilization: adaptation across biologically relevant ‘levels’ in tamarins. BMC ecology and evolution, 24(1), 22. 10.1186/s12862-023-02193-z.

Berry, J. L., Towers, J. D., Webber, R. L., Pope, T. L., Davidai, G., & Zimmerman, M. (1996). Change in trabecular architecture as measured by fractal dimension. Journal of biomechanics, 29(6), 819–822. 10.1016/0021-9290(95)00113-1.

Biewener, A.A., Fazzalari, N.L., Konieczynski, D.D., Baudinette, R.V., 1996. Adaptive changes in trabecular architecture in relation to functional strain patterns and disuse. Bone 19, 1–8. 10.1016/8756-3282(96)00116-0

Bishop, P. J., Hocknull, S. A., Clemente, C. J., Hutchinson, J. R., Farke, A. A., Beck, B. R., Barrett, R. S., & Lloyd, D. G. (2018). Cancellous bone and theropod dinosaur locomotion. Part I—an examination of cancellous bone architecture in the hindlimb bones of theropods. PeerJ, 6, e5778. 10.7717/peerj.5778.

Bourne, M., Sinkler, M. A., & Murphy, P. B. (2018). Anatomy, bony pelvis and lower limb, tibia.

Boyer, D. M., Toussaint, S., & Godinot, M. (2017). Postcrania of the most primitive euprimate and implications for primate origins. Journal of Human Evolution, 111, 202–215. 10.1016/j.jhevol.2017.07.005.

Carlson, K.J., Lublinsky, S., Judex, S., 2008. Do different locomotor modes during growth modulate trabecular architecture in the murine hind limb? Integrative and Comparative Biology 48, 385–393. 10.1093/icb/icn066

Cendre, E., Mitton, D., Roux, J.P., Arlot, M.E., Duboeuf, F., Burt-Pichat, B. et al. (1999) High-resolution computed tomography for architectural characterization of human lumbar cancellous bone: relationships with histomorphometry and biomechanics. Osteoporosis International, 10, 353–360.

Chang, G., Pakin, S. K., Schweitzer, M. E., Saha, P. K., & Regatte, R. R. (2008). Adaptations in trabecular bone microarchitecture in Olympic athletes determined by 7T MRI. Journal of Magnetic Resonance Imaging: An Official Journal of the International Society for Magnetic Resonance in Medicine, 27(5), 1089–1095. 10.1002/jmri.21326.

Channon, A. J., Crompton, R. H., Günther, M. M., D’août, K., & Vereecke, E. E. (2010). The biomechanics of leaping in gibbons. American Journal of Physical Anthropology, 143(3), 403–416. 10.1002/ajpa.21329.

Chappard, D., Chennebault, A., Moreau, M., Legrand, E., Audran, M., & Basle, M. F. (2001). Texture analysis of X-ray radiographs is a more reliable descriptor of bone loss than mineral content in a rat model of localized disuse induced by the *Clostridium botulinum* toxin. Bone, 28(1), 72–79. 10.1016/S8756-3282(00)00438-5.

Chirchir, H., Zeininger, A., Nakatsukasa, M., Ketcham, R. A., & Richmond, B. G. (2017). Does trabecular bone structure within the metacarpal heads of primates vary with hand posture?. Comptes Rendus Palevol, 16(5-6), 533–544. 10.1016/j.crpv.2016.10.002.

Connour, J. R., Glander, K., & Vincent, F. (2000). Postcranial adaptations for leaping in primates. Journal of Zoology, 251(1), 79–103.

Crane, A. H., Baldry, C. J., Rankin, K. E., Clarkin, C. E., Williams, K. A., & Gostling, N. J. (2025). The three-dimensional structure of medullary bone: Novel criteria for the identification of avian sex-specific bone tissue. Developmental Biology, 521, 108–121. 10.1016/j.ydbio.2025.02.007.

Crompton, R. H., Blanchard, M. L., Coward, S., Alexander, R. M., & Thorpe, S. K. (2010). Vertical clinging and leaping revisited: locomotion and habitat use in the western tarsier, Tarsius bancanus explored via loglinear modeling. International journal of primatology, 31, 958–979.

Deckers, K., Tsegai, Z. J., Skinner, M. M., Zeininger, A., & Kivell, T. L. (2022). Ontogenetic changes to metacarpal trabecular bone structure in mountain and western lowland gorillas. Journal of Anatomy, 241(1), 82–100. 10.1111/joa.13630.

Demes, B., Fleagle, J. G., & Jungers, W. L. (1999). Takeoff and landing forces of leaping strepsirhine primates. Journal of Human Evolution, 37(2), 279–292. 10.1006/jhev.1999.0311.

Demes, B., Jungers, W. L., Fleagle, J. G., Wunderlich, R. E., Richmond, B. G., & Lemelin, P. (1996). Body size and leaping kinematics in Malagasy vertical clingers and leapers. Journal of Human Evolution, 31(4), 367–388.

Demes, B., Jungers, W. L., Gross, T. S., & Fleagle, J. G. (1995). Kinetics of leaping primates: influence of substrate orientation and compliance. American Journal of Physical Anthropology, 96(4), 419–429. 10.1002/ajpa.1330960407.

Doube, M., Kłosowski, M. M., Arganda-Carreras, I., Cordelières, F. P., Dougherty, R. P., Jackson, J. S., … & Shefelbine, S. J. (2010). BoneJ: free and extensible bone image analysis in ImageJ. Bone, 47(6), 1076–1079.

Doube, M., Kłosowski, M. M., Wiktorowicz-Conroy, A. M., Hutchinson, J. R., & Shefelbine, S. J. (2011). Trabecular bone scales allometrically in mammals and birds. Proceedings of the Royal Society B: Biological Sciences, 278(1721), 3067–3073. 10.1098/rspb.2011.0069.

Dunmore, C. J., Bachmann, S., Synek, A., Pahr, D. H., Skinner, M. M., & Kivell, T. L. (2024). The deep trabecular structure of first metacarpals in extant hominids. American Journal of Biological Anthropology, 183(3), e24695. 10.1002/ajpa.24695.

Emerson, S. B. (1978). Allometry and jumping in frogs: helping the twain to meet. Evolution, 551–564.

Fajardo, R. J., & Müller, R. (2001). More on the three-dimensional trabecular architecture of anthropoid primates: *Macaca fascicularis* and *Symphalangus syndactylus*. Am. J. Phys. Anthropol.

Fajardo, R. J., Müller, R., Ketcham, R. A., & Colbert, M. (2007). Nonhuman anthropoid primate femoral neck trabecular architecture and its relationship to locomotor mode. The Anatomical Record: Advances in Integrative Anatomy and Evolutionary Biology: Advances in Integrative Anatomy and Evolutionary Biology, 290(4), 422–436. 10.1002/ar.20493. Feder, J. (1988) Fractals. New York: Plenum.

Feltrin, G. P., Stramare, R., Miotto, D., Giacomini, D., & Saccavini, C. (2004). Bone fractal analysis. Current Osteoporosis Reports, 2(2), 53–58. 10.1007/s11914-004-0004-4.

Fyhrie, D. P., & Zauel, R. (2015). Directional tortuosity as a predictor of modulus damage for vertebral cancellous bone. Journal of biomechanical engineering, 137(1), 011007. 10.1115/1.4029177.

Garber, P. A. (1991). A comparative study of positional behavior in three species of tamarin monkeys. Primates, 32, 219–230. 10.1007/BF02381179.

Garber, P. A., & Pruetz, J. D. (1995). Positional behavior in moustached tamarin monkeys: effects of habitat on locomotor variability and locomotor stability. Journal of Human Evolution, 28(5), 411–426. 10.1006/jhev.1995.1032.

Garber, P. A., McKenney, A. C., & Mallott, E. K. (2012). The ecology of trunk-to-trunk leaping in *Saguinus fuscicollis*: implications for understanding locomotor diversity in Callitrichines. Neotropical Primates, 19(1), 1–7. 10.1896/044.019.0101.

Garber, P. A., & Teaford, M. F. (1986). Body weights in mixed species troops of Saguinus mystax mystax and Saguinus fuscicollis nigrifrons in Amazonian Peru. American Journal of Physical Anthropology, 71(3), 331–336. 10.1002/ajpa.1330710308.

Georgiou, L., Kivell, T.L., Pahr, D.H., Buck, L.T., Skinner, M.M., 2019. Trabecular architecture of the great ape and human femoral head. J. Anat. 234, 679–693. 10.1111/joa.12957

Geraets, W. G., & Van Der Stelt, P. F. (2000). Fractal properties of bone. Dentomaxillofacial Radiology, 29(3), 144–153. 10.1038/sj/dmfr/4600524.

Gônet, J., Laurin, M., Hutchinson, J.R., 2023. Evolution of posture in amniotes– Diving into the trabecular architecture of the femoral head. Journal of Evolutionary Biology 36(8), 1150–1165.

Harper, C. M., & Patel, B. A. (2024). Trabecular bone variation in the gorilla calcaneus. American Journal of Biological Anthropology, 184(3), e24939. 10.1002/ajpa.24939.

Harrigan TP, Mann RW (1984) Characterization of microstructural anisotropy in orthotropic materials using a second rank tensor. J Mater Sci 19, 761–767.

Havill, L. M., Allen, M. R., Bredbenner, T. L., Burr, D. B., Nicolella, D. P., Turner, C. H., Warren, D. M., & Mahaney, M. C. (2010). Heritability of lumbar trabecular bone mechanical properties in baboons. Bone, 46(3), 835–840. 10.1016/j.bone.2009.11.002.

Huiskes, R., Ruimerman, R., Van Lenthe, G. H., & Janssen, J. D. (2000). Effects of mechanical forces on maintenance and adaptation of form in trabecular bone. Nature, 405(6787), 704–706. 10.1038/35015116.

Ingle, D. N., & Porter, M. E. (2021). Microarchitecture of cetacean vertebral trabecular bone among swimming modes and diving behaviors. Journal of Anatomy, 238(3), 643–652.

Garber, P. A., & Teaford, M. F. (1986). Body weights in mixed species troops of Saguinus mystax mystax and Saguinus fuscicollis nigrifrons in Amazonian Peru. American Journal of Physical Anthropology, 71(3), 331–336. 10.1111/joa.13329.

Ingle, D. N., & Porter, M. E. (2021). Microarchitecture of cetacean vertebral trabecular bone among swimming modes and diving behaviors. Journal of Anatomy, 238(3), 643–652. 10.1111/joa.13329.

Iura, A., McNerny, E. G., Zhang, Y., Kamiya, N., Tantillo, M., Lynch, M., Kohn, D. H. & Mishina, Y. (2015). Mechanical loading synergistically increases trabecular bone volume and improves mechanical properties in the mouse when BMP signaling is specifically ablated in osteoblasts. PloS one, 10(10), e0141345. 10.1371/journal.pone.0141345.

Lazenby, R. A., Skinner, M. M., Kivell, T. L., & Hublin, J. J. (2011). Scaling VOI size in 3D _μ_CT studies of trabecular bone: A test of the over_-_sampling hypothesis. American Journal of Physical Anthropology, 144(2), 196–203.

Jacquet, G., Ohley, W. J., Mont, M. A., Siffert, R., & Schmukler, R. (1990). Measurement of bone structure by use of fractal dimension. In [1990] Proceedings of the Twelfth Annual International Conference of the IEEE Engineering in Medicine and Biology Society (pp. 1402–1403). IEEE. 10.1109/IEMBS.1990.69181.

James, R. S., & Wilson, R. S. (2008). Explosive jumping: extreme morphological and physiological specializations of Australian rocket frogs (Litoria nasuta). Physiological and Biochemical Zoology, 81(2), 176–185.

Keaveny, T. M., Morgan, E. F., Niebur, G. L., & Yeh, O. C. (2001). Biomechanics of trabecular bone. Annual review of biomedical engineering, 3(1), 307–333. 10.1146/annurev.bioeng.3.1.307.

Kivell, T. L., Skinner, M. M., Lazenby, R., & Hublin, J. J. (2011). Methodological considerations for analyzing trabecular architecture: an example from the primate hand. Journal of anatomy, 218(2), 209–225.

Kivell, T. L. (2016). A review of trabecular bone functional adaptation: what have we learned from trabecular analyses in extant hominoids and what can we apply to fossils?. Journal of Anatomy, 228(4), 569–594. 10.1111/joa.12446.

Kivell, T.L., Davenport, R., Hublin, J.-J., Thackeray, J.F., Skinner, M.M., 2018. Trabecular architecture and joint loading of the proximal humerus in extant hominoids, Ateles , and Australopithecus africanus. Am J Phys Anthropol 167, 348–365. 10.1002/ajpa.23635

Lukova, A., Dunmore, C. J., Bachmann, S., Synek, A., Pahr, D. H., Kivell, T. L., & Skinner, M. M. (2024). Trabecular architecture of the distal femur in extant hominids. Journal of Anatomy, 245(1), 156–180. 10.1111/joa.14026.

Majumdar, S., Genant, H. K., Grampp, S., Newitt, D. C., Truong, V. H., Lin, J. C., & Mathur, A. (1997). Correlation of trabecular bone structure with age, bone mineral density, and osteoporotic status: in vivo studies in the distal radius using high resolution magnetic resonance imaging. Journal of Bone and Mineral Research, 12(1), 111–118. 10.1359/jbmr.1997.12.1.111.

Maquer, G., Musy, S.N., Wandel, J., Gross, T., Zysset, P.K., 2015. Bone volume fraction and fabric anisotropy are better determinants of trabecular bone stiffness than other morphological variables. J. Bone Miner. Res. 30, 1000–1008. 10.1002/jbmr.2437

Messent, E. A., Buckland-Wright, J. C., & Blake, G. M. (2005a). Fractal analysis of trabecular bone in knee osteoarthritis (OA) is a more sensitive marker of disease status than bone mineral density (BMD). Calcified tissue international, 76, 419–425. 10.1007/s00223-004-0160-7.

Messent, E. A., Ward, R. J., Tonkin, C. J., & Buckland-Wright, C. (2005b). Tibial cancellous bone changes in patients with knee osteoarthritis. A short-term longitudinal study using Fractal Signature Analysis. Osteoarthritis and cartilage, 13(6), 463–470. 10.1016/j.joca.2005.01.007.

Mielke, M., Wölfer, J., Arnold, P., van Heteren, A. H., Amson, E., & Nyakatura, J. A. (2018). Trabecular architecture in the sciuromorph femoral head: allometry and functional adaptation. Zoological letters, 4, 1–11. 10.1186/s40851-018-0093-z.

Mo, X., Ge, W., Miraglia, M., Inglese, F., Zhao, D., Stefanini, C., & Romano, D. (2020). Jumping locomotion strategies: From animals to bioinspired robots. Applied Sciences, 10(23), 8607.

Modlesky, C. M., Subramanian, P., & Miller, F. (2008). Underdeveloped trabecular bone microarchitecture is detected in children with cerebral palsy using high-resolution magnetic resonance imaging. Osteoporosis international, 19, 169–176. 10.1007/s00198-007-0433-x.

Nauwelaerts, S., & Aerts, P. (2006). Take-off and landing forces in jumping frogs. Journal of Experimental Biology, 209(1), 66–77. 10.1242/jeb.01969.

Nguyen, U., Alfieri, F., Veneziano, A., Licht, A., Nyakatura J.A., (2025). Data from: Trabecular structure correlates with leaping distance in tamarins. https://figshare.com/s/e803d685895a41ec9f57.

Nyakatura, J. A., & Heymann, E. W. (2010). Effects of support size and orientation on symmetric gaits in free-ranging tamarins of Amazonian Peru: implications for the functional significance of primate gait sequence patterns. Journal of Human Evolution, 58(3), 242–251. 10.1016/j.jhevol.2009.11.010.

Parkinson, I. H., & Fazzalari, N. L. (2012). Characterisation of trabecular bone structure. In Skeletal aging and osteoporosis: biomechanics and mechanobiology (pp. 31–51). Berlin, Heidelberg: Springer Berlin Heidelberg. 10.1007/8415_2011_113.

Parkinson, I. H., Badiei, A., Stauber, M., Codrington, J., Müller, R., & Fazzalari, N. L. (2012). Vertebral body bone strength: the contribution of individual trabecular element morphology. Osteoporosis International, 23, 1957–1965. 10.1007/s00198-011-1832-6.

Paternoster, L., Lorentzon, M., Lehtimäki, T., Eriksson, J., Kähönen, M., Raitakari, O., … & Ohlsson, C. (2013). Genetic determinants of trabecular and cortical volumetric bone mineral densities and bone microstructure. PLoS genetics, 9(2), e1003247. 10.1371/journal.pgen.1003247.

Pau, G., Fuchs, F., Sklyar, O., Boutros, M., & Huber, W. (2010). EBImage-an R package for image processing with applications to cellular phenotypes. Bioinformatics, 26(7), 979–981. 10.1093/bioinformatics/btq046.

Pontzer, H., Lieberman, D. E., Momin, E., Devlin, M. J., Polk, J. D., Hallgrimsson, B., & Cooper, D. M. L. (2006). Trabecular bone in the bird knee responds with high sensitivity to changes in load orientation. Journal of Experimental Biology, 209(1), 57–65. 10.1242/jeb.01971.

Polvadore, T., McGraw, W. S., & Daegling, D. J. (2024). Limb and hip morphology of two African colobine monkeys and its relationship to the mechanics of leaping and bounding locomotion. American Journal of Biological Anthropology, 183(1), 92–106.

Pornprasertsuk, S., Ludlow, J. B., Webber, R. L., Tyndall, D. A., Sanhueza, A. I., & Yamauchi, M. (2001). Fractal dimension analysis of weight-bearing bones of rats during skeletal unloading. Bone, 29(2), 180–184. 10.1016/S8756-3282(01)00493-8.

Preuschoft, H., Hohn, B., Scherf, H., Schmidt, M., Krause, C., & Witzel, U. (2010). Functional analysis of the primate shoulder. International journal of primatology, 31(2), 301–320.

R Core Team (2023). R: A language and environment for statistical computing. R Foundation for Statistical Computing, Vienna, Austria. https://www.R-project.org/.

Ragni, A. J. (2020). Trabecular architecture of the capitate and third metacarpal through ontogeny in chimpanzees (*Pan troglodytes*) and gorillas (*Gorilla gorilla*). Journal of Human Evolution, 138, 102702.

Roque, W. L., & Alberich-Bayarri, A. (2015). Tortuosity influence on the trabecular bone elasticity and mechanical competence. Developments in medical image processing and computational vision, 173–191. 10.1007/978-3-319-13407-9_1.

Roque, W. L., Arcaro, K., & Lanfredi, R. B. (2012). Trabecular network tortuosity and connectivity of distal radius from microtomographic images. Revista Brasileira de Engenharia Biomédica, 28, 116–123. 10.4322/rbeb.2012.017.

Ruff, C. B. (2002). Long bone articular and diaphyseal structure in Old World monkeys and apes. I: locomotor effects. American Journal of Physical Anthropology: The Official Publication of the American Association of Physical Anthropologists, 119(4), 305–342. 10.1002/ajpa.10117.

Ryan, T. M., & Ketcham, R. A. (2001). Femoral head trabecular bone structure in two omomyid primates. Journal of Human Evolution, 43(2), 241–263. 10.1006/jhev.2002.0575.

Ryan, T. M., & Ketcham, R. A. (2002). The three-dimensional structure of trabecular bone in the femoral head of strepsirrhine primates. Journal of human evolution, 43(1), 1–26. 10.1006/jhev.2002.0552.

Ryan, T. M., & Ketcham, R. A. (2005). Angular orientation of trabecular bone in the femoral head and its relationship to hip joint loads in leaping primates. Journal of Morphology, 265(3), 249–263. 10.1002/jmor.10315.

Ryan, T. M., & Shaw, C. N. (2012). Unique suites of trabecular bone features characterize locomotor behavior in human and non-human anthropoid primates. PloS one, 7(7), e41037. 10.1371/journal.pone.0041037.

Ryan, T. M., & Shaw, C. N. (2015). Gracility of the modern Homo sapiens skeleton is the result of decreased biomechanical loading. Proceedings of the National Academy of Sciences, 112(2), 372–377. 10.1073/pnas.1418646112.

Ryan, T. M., & Van Rietbergen, B. (2005). Mechanical significance of femoral head trabecular bone structure in Loris and Galago evaluated using micromechanical finite element models. American Journal of Physical Anthropology: The Official Publication of the American Association of Physical Anthropologists, 126(1), 82–96. 10.1002/ajpa.10414.

Ryan, T. M., & Walker, A. (2010). Trabecular bone structure in the humeral and femoral heads of anthropoid primates. The Anatomical Record: Advances in Integrative Anatomy and Evolutionary Biology, 293(4), 719–729. 10.1002/ar.21139.

Ryan, T. M., Colbert, M., Ketcham, R. A., & Vinyard, C. J. (2010). Trabecular bone structure in the mandibular condyles of gouging and nongouging platyrrhine primates. American Journal of Physical Anthropology: The Official Publication of the American Association of Physical Anthropologists, 141(4), 583–593. 10.1002/ajpa.21178.

Saers, J. P. P. (2017). Ontogeny and functional adaptation of trabecular bone in the human foot (Doctoral dissertation). 10.17863/CAM.17149.

Saers, J. P. P., Gordon, A. D., Ryan, T. M., & Stock, J. T. (2022). Growth and development of trabecular structure in the calcaneus of Japanese macaques (*Macaca fuscata*) reflects locomotor behavior, life history, and neuromuscular development. Journal of Anatomy, 241(1), 67–81. 10.1111/joa.13641.

Saparin, P., Scherf, H., Hublin, J. J., Fratzl, P., & Weinkamer, R. (2011). Structural adaptation of trabecular bone revealed by position resolved analysis of proximal femora of different primates. The Anatomical Record: Advances in Integrative Anatomy and Evolutionary Biology, 294(1), 55–67. 10.1002/ar.21285.

Schneider, C. A., Rasband, W. S., & Eliceiri, K. W. (2012). NIH Image to ImageJ: 25 years of image analysis. Nature methods, 9(7), 671–675. 10.1038/nmeth.2089.

Smith, S. M., & Angielczyk, K. D. (2020). Deciphering an extreme morphology: bone microarchitecture of the hero shrew backbone (Soricidae: *Scutisorex*). Proceedings of the Royal Society B, 287(1926), 20200457. 10.1098/rspb.2020.0457.

Southard, T. E., Southard, K. A., Jakobsen, J. R., Hillis, S. L., & Najim, C. A. (1996). Fractal dimension in radiographic analysis of alveolar process bone. Oral Surgery, Oral Medicine, Oral Pathology, Oral Radiology, and Endodontology, 82(5), 569–576. 10.1016/S1079-2104(96)80205-8.

Sowińska-Przepiera, E., Krzyścin, M., Syrenicz, I., Ćwiertnia, A., Orlińska, A., Ćwiek, D., Branecka-Woźniak, D., Cymbaluk-Płoska, A., Bumbulienė, Z. & Syrenicz, A. (2024). Evaluation of Trabecular Bone Microarchitecture and Bone Mineral Density in Young Women, Including Selected Hormonal Parameters. Biomedicines, 12(4), 758. 10.3390/biomedicines12040758.

Stauber, M., Rapillard, L., van Lenthe, G.H., Zysset, P., Müller, R., 2006. Importance of individual rods and plates in the assessment of bone quality and their contribution to bone stiffness. Journal of Bone and Mineral Research 21(4), 586–595.

Sugiyama, T., Price, J. S., & Lanyon, L. E. (2010). Functional adaptation to mechanical loading in both cortical and cancellous bone is controlled locally and is confined to the loaded bones. Bone, 46(2), 314–321. 10.1016/j.bone.2009.08.054.

Terranova, C. J. (1995). Leaping behaviors and the functional morphology of strepsirhine primate long bones. Folia Primatologica, 65(4), 181–201.

Tsegai, Z. J., Kivell, T. L., Gross, T., Nguyen, N. H., Pahr, D. H., Smaers, J. B., & Skinner, M. M. (2013). Trabecular bone structure correlates with hand posture and use in hominoids. PLoS One, 8(11), e78781. 10.1371/journal.pone.0078781.

Tsegai, Z. J., Skinner, M. M., Pahr, D. H., Hublin, J. J., & Kivell, T. L. (2018). Ontogeny and variability of trabecular bone in the chimpanzee humerus, femur and tibia. American Journal of Physical Anthropology, 167(4), 713–736. 10.1002/ajpa.23696.

Turner, C. H., Hsieh, Y. F., Müller, R., Bouxsein, M. L., Baylink, D. J., Rosen, C. J., Grynpas, M.D., Donahue, L.R., & Beamer, W. G. (2000). Genetic regulation of cortical and trabecular bone strength and microstructure in inbred strains of mice. Journal of Bone and Mineral Research, 15(6), 1126–1131. 10.1359/jbmr.2000.15.6.1126.

Veneziano, A., Cazenave, M., Alfieri, F., Panetta, D., & Marchi, D. (2021). Novel strategies for the characterization of cancellous bone morphology: Virtual isolation and analysis. American Journal of Physical Anthropology, 175(4), 920–930. 10.1002/ajpa.24272.

Wallace, I. J., Tommasini, S. M., Judex, S., Garland Jr, T., & Demes, B. (2012). Genetic variations and physical activity as determinants of limb bone morphology: an experimental approach using a mouse model. American Journal of Physical Anthropology, 148(1), 24–35. 10.1002/ajpa.22028.

Weatherholt, A. M., Fuchs, R. K., & Warden, S. J. (2013). Cortical and trabecular bone adaptation to incremental load magnitudes using the mouse tibial axial compression loading model. Bone, 52(1), 372–379. 10.1016/j.bone.2012.10.026.

Zack, E. H., Smith, S. M., & Angielczyk, K. D. (2022). Effect of captivity on the vertebral bone microstructure of xenarthran mammals. The Anatomical Record, 305(7), 1611–1628.

