## Supplementary Material for "Trabecular structure correlates with leaping distance in tamarins"

[5]: Archéozoologie, Archéobotanique: Sociétés, Pratiques et Environnements (AASPE), Muséum National d'Histoire Naturelle, CNRS, Paris, France

[+]:

[*]: These authors contributed equally.

#### Supplementary Material

**Note S1:** As further detailed in Veneziano et al. 2021 and Alfieri et al. 2025, the four topological indices that we quantified on whole epiphyses inform on structural properties that, in turn, have recognised biomechanical meaning

1. **Node density:** this index derives from trabecular density which is, in turn, a conceptual generalization of BV/TV. For instance, Tsegai et al. 2018 have computed BV/TV exploiting a grid which encompassed the bone and then used an interpolation to extract a continuous BV/TV map. If TV is replaced with a unit of volume, the same procedure results in a density map. If, in this procedure, we add the skeletonisation procedure (see Methods, in the main text), then the density distribution only includes topological properties and we then obtain a node density distribution. Node density informs on the distribution of the number of nodes within the volume (Veneziano et al. 2021) and it was computed by estimating a kernel-density over a 3D grid that is the whole-epiphysis in this study (TV). Hence, since node density is indirectly related to BV/TV, we can assume that they similarly respond to biomechanical stimuli - i.e. positively relating to increased loadings - as it was suggested by preliminary results on the femoral head of humans, African apes, quadrupeds and brachiators (Veneziano et al. 2021). To further support the positive relationship between node density and increased loadings, Cendre et al. (1999) found that node density/total volume increases in association with higher maximum compressive strenght.
2. **Trabecular length:** After the procedure of skeletonization, it is possible to compute the trabecular length, which is the geodesic length of a branch, delimited by a couple of nodes. The potential functional meaning of trabecular lenght lies in its expected negative correlation with trabecular densit, i.e. denser structures are generally characterized by shorter trabeculae. Moreover, the trabecular length itself may be related to bone strength (Parkinson et al. 2012) and we can thus assume that shorter trabeculae can be expected in association with denser structures in response to increased loadings.
3. **Trabecular tortuosity:** the computation of trabecular length is needed to compute trabecular tortuosity, a proxy for how trabeculae tend to be curved and convoluted. If the trabecular length between two nodes (computed as detailed above) is divided by the euclidean length between the same two nodes, we obtain the trabecular tortuosity. In other words, if a given arc length

occurs over a shorter euclidean distance, it means that the branch (hence, the trabecula) is more sinuous, and it contributes to make the whole structure more tortuous. Tortuosity is a property that has been shown to capture mechanical features (Fyhrie and Zauel 2015). Indeed, tortuosity affects how flexibly the trabeculae respond to mechanical loads: higher tortuosity correlates with lower stiffness, i.e. higher elasticity (Roque et al. 2012; Roque and Alberich-Bayarri 2015). If the stiffness decreases, and then the tortuosity and flexibility increase, it means than the entire structure is more elastic, and it may suggest a response to higher loadings. Hence, tortuosity can positively correlate with loadings.

4. **Fractal dimension:** it measures the structural complexity, and specifically how a structure tends to be similar to itself when the degree of magnification changes. The more the structure resembles itself at varying degrees of magnification, the more the structure approaches an ideal fractal network (Feder 1988; Pornprasertsuk et al. 2001). This concept has tbeen applied to trabecular structure to study mechanical aspects related to fracture risk and osseous pathologies in humans (Feltrin et al. 2004; Messent et al. 2005b,a) and to bone density, that informs on bone strenght (Ammann and Rizzoli 2003). FD relates to density and we here assume that they correlate negatively, based on experimental evidence, i.e. if bone is experimentally forced to experience periods of inactivity, it responds by decreasing density and increase FD (Pornprasertsuk et al. 2001). Accordingly, we expect FD to decrease in association with increased loadings.

**Table S1:** Bone lengths and scanning information of the twelve analyzed humeri/tibiae

| Species | Specimen | Acquired at | CT-scanning resolution (mm) | CT scanning Voltage (kV) | CT scanning current (uA) | Filter | Humerus length (mm) | Tibia length (mm) |
|---|---|---|---|---|---|---|---|---|
| *L.nigrifrons* | FMNH 122268 | UChicago | 0.0165 | 100 | 165 | 0.2mm Cu | 52.338 | 70.422 |
| *L.nigrifrons* | FMNH 122269 | UChicago | 0.0165 | 100 | 165 | 0.2mm Cu | 52.420 | 66.478 |
| *L.nigrifrons* | FMNH 122270 | UChicago | 0.0165 | 100 | 165 | 0.2mm Cu | 51.348 | 66.709 |
| *S.mystax* | AMNH 188173 | SMiF | 0.015487 | 170 | 79 | No | 49.745 | 63.745 |
| *S.mystax* | AMNH 188177 | SMiF | 0.015487 | 172 | 85 | No | 46.895 | 59.641 |
| *S.mystax* | AMNH 188178 | SMiF | 0.015487 | 170 | 83 | No | 50.287 | 64.426 |
| *S.imperator* | FMNH 98035 | UChicago | 0.0165 | 100 | 165 | 0.2mm Cu | 53.542 | 67.188 |
| *S.imperator* | FMNH 98036 | UChicago | 0.0165 | 100 | 165 | 0.2mm Cu | 53.592 | 66.165 |
| *S.imperator* | FMNH 121551 | UChicago | 0.018 | 100 | 165 | 0.2mm Cu | 55.836 | 70.884 |
| *S.midas* | FMNH 93239 | UChicago | 0.0165 | 100 | 165 | 0.2mm Cu | 57.139 | - |
| *S.midas* | FMNH 93236 | UChicago | 0.0165 | 100 | 165 | 0.2mm Cu | 55.209 | - |
| *S.midas* | FMNH 93516 | UChicago | 0.0165 | 100 | 165 | 0.2mm Cu | 56.925 | 75.438 |

**Table S2:** Raw results for the topological indices and traditional trabecular variables extracted from humeral epiphyses

Humeral proximal epiphyses

| Species | Specimen | NodDen_mean | NodDen_median | TrabLen | TrabTort_mean | TrabTort_median | FD | DA | BV/TV |
|---|---|---|---|---|---|---|---|---|---|
| L. nigrifrons | FMNH_122268 | 0.01689643 | 0.01653007 | 0.21250315 | 1.12889552 | 1.06037447 | 2.04154152 | 0.2994692019 | 0.0850497933 |
| L. nigrifrons | FMNH_122269 | 0.01516662 | 0.01372665 | 0.21193715 | 1.13252471 | 1.06231243 | 2.13937887 | 0.2336497064 | 0.1710026627 |
| L. nigrifrons | FMNH_122270 | 0.01364965 | 0.01293283 | 0.19470246 | 1.11614431 | 1.05726686 | 2.18581409 | 0.3036563496 | 0.1326528135 |
| S.mystax | AMNH_188173 | 0.01526758 | 0.01440248 | 0.22393557 | 1.1231462 | 1.06035569 | 2.08736926 | 0.2699711709 | 0.190463774 |
| S.mystax | AMNH_188177 | 0.01725113 | 0.01693791 | 0.22384094 | 1.11809448 | 1.06061091 | 2.10629236 | 0.4014398649 | 0.1848945299 |
| S.mystax | AMNH_188178 | 0.01602279 | 0.01556561 | 0.2259206 | 1.12635744 | 1.06026407 | 2.08776903 | 0.2915342123 | 0.196831586 |
| S.imperator | FMNH_98036 | 0.00923939 | 0.00911521 | 0.21936717 | 1.11434779 | 1.05968505 | 2.17469472 | 0.2967587141 | 0.1762331189 |
| S.imperator | FMNH_121551 | 0.00893315 | 0.00892911 | 0.20300627 | 1.11092799 | 1.06046037 | 2.29642146 | 0.3127146776 | 0.189336165 |
| S.midas | FMNH_93239 | 0.01095205 | 0.01038249 | 0.2373098 | 1.11644196 | 1.05808405 | 2.0485198 | 0.2774638209 | 0.1534299294 |
| S.midas | FMNH_93516 | 0.00788591 | 0.00751383 | 0.22221041 | 1.11589279 | 1.05718546 | 2.15454963 | 0.286950079 | 0.1920903086 |

Humeral distal epiphyses

| Species | Specimen | NodDen_mean | NodDen_median | TrabLen | TrabTort_mean | TrabTort_median | FD | DA | BV/TV |
|---|---|---|---|---|---|---|---|---|---|
| L.nigrifrons | FMNH_122268 | 0.0352776 | 0.032986 | 0.18374689 | 1.15664737 | 1.06300132 | 2.06300405 | 0.3536145576 | 0.1616580373 |
| L.nigrifrons | FMNH_122269 | 0.05989379 | 0.05543621 | 0.18433095 | 1.17358021 | 1.06837196 | 2.0327086 | 0.5186126949 | 0.23047478 |
| L.nigrifrons | FMNH_122270 | 0.05170705 | 0.04914879 | 0.18342762 | 1.14890554 | 1.06089379 | 2.14386731 | 0.4818634811 | 0.2290525036 |
| S.mystax | AMNH_188173 | 0.0680265 | 0.06518007 | 0.2092837 | 1.16592152 | 1.07622962 | 1.78149961 | 0.521837972 | 0.1672183554 |
| S.mystax | AMNH_188177 | 0.07973235 | 0.07293333 | 0.20254803 | 1.17013673 | 1.06862335 | 1.85222329 | 0.4892486069 | 0.1889769505 |
| S.mystax | AMNH_188178 | 0.08653515 | 0.0832564 | 0.20318213 | 1.19017183 | 1.06832703 | 1.7064093 | 0.4844072018 | 0.1558758403 |
| S.imperator | FMNH_98035 | 0.03672757 | 0.03410344 | 0.1728478 | 1.14182806 | 1.05870741 | 2.16893836 | 0.4793085966 | 0.2842036962 |
| S.imperator | FMNH_98036 | 0.02766595 | 0.02674902 | 0.20570805 | 1.13358862 | 1.0624972 | 2.12157108 | 0.3741943257 | 0.240996279 |
| S.imperator | FMNH_121551 | 0.0235465 | 0.02276308 | 0.1921667 | 1.12556552 | 1.0624795 | 2.19677409 | 0.3128088608 | 0.2421827378 |
| S.midas | FMNH_93239 | 0.02680486 | 0.02575404 | 0.21526579 | 1.15195741 | 1.06379119 | 2.01214074 | 0.4859181483 | 0.2038745845 |
| S.midas | FMNH_93236 | 0.02232845 | 0.02076193 | 0.17472116 | 1.14034822 | 1.05969714 | 2.16775709 | 0.5091363675 | 0.2458056698 |
| S.midas | FMNH_93516 | 0.02628588 | 0.02477753 | 0.20956465 | 1.14872888 | 1.06317054 | 2.00017572 | 0.5054128506 | 0.203715211 |

**Table S3:** Raw results for the topological indices and traditional trabecular variables extracted from tibial epiphyses

Tibial proximal epiphyses

| Species | Specimen | NodDen_mean | NodDen_median | TrabLen | TrabTort_mean | TrabTort_median | FD | DA | BV/TV |
|---|---|---|---|---|---|---|---|---|---|
| L. nigrifrons | FMNH_122268 | 0.01797327 | 0.01480379 | 0.17531275 | 1.14804527 | 1.06579176 | 2.09205693 | 0.3199313162 | 0.1313727278 |
| L. nigrifrons | FMNH_122269 | 0.01409536 | 0.01292352 | 0.20635195 | 1.12730837 | 1.06052408 | 2.1873939 | 0.3184675498 | 0.2335995473 |
| L. nigrifrons | FMNH_122270 | 0.0143671 | 0.01363609 | 0.18962899 | 1.1102672 | 1.05705751 | 2.16318145 | 0.3086971957 | 0.1456952133 |
| S. mystax | AMNH_188173 | 0.01441196 | 0.01407099 | 0.21777114 | 1.11769237 | 1.06015572 | 2.15681309 | 0.2581519409 | 0.2228000773 |
| S. mystax | AMNH_188177 | 0.01815416 | 0.01690632 | 0.22906937 | 1.11233111 | 1.06197188 | 2.0827739 | 0.4703001606 | 0.1949843215 |
| S. mystax | AMNH_188178 | 0.01528028 | 0.01421273 | 0.2182397 | 1.12916652 | 1.0621106 | 2.11240575 | 0.3470630233 | 0.2366615044 |
| S. imperator | FMNH_98035 | 0.01717169 | 0.0147536 | 0.14977164 | 1.09897389 | 1.05131301 | 2.30187632 | 0.2494661986 | 0.2625664815 |
| S. imperator | FMNH_98036 | 0.01081666 | 0.01060486 | 0.21948656 | 1.11141683 | 1.0608389 | 2.18448485 | 0.3223542523 | 0.1975674889 |
| S. imperator | FMNH_121551 | 0.01015415 | 0.00984192 | 0.20363324 | 1.11686282 | 1.05958448 | 2.19641234 | 0.3669707048 | 0.2047152717 |
| S. midas | FMNH_93516 | 0.00725621 | 0.00664052 | 0.20479126 | 1.11314952 | 1.05663358 | 2.20665397 | 0.3201412069 | 0.2180968338 |

Tibial distal epiphyses

| Species | Specimen | NodDen_mean | NodDen_median | TrabLen | TrabTort_mean | TrabTort_median | FD | DA | BV/TV |
|---|---|---|---|---|---|---|---|---|---|
| L. nigrifrons | FMNH_122268 | 0.03457818 | 0.03177105 | 0.18059852 | 1.12979591 | 1.0597556 | 2.07597361 | 0.1976391448 | 0.121813684 |
| L. nigrifrons | FMNH_122269 | 0.03979007 | 0.03653906 | 0.18737309 | 1.1367426 | 1.06066531 | 2.04143224 | 0.2439748075 | 0.1964030982 |
| L. nigrifrons | FMNH_122270 | 0.04239635 | 0.03984305 | 0.18101821 | 1.12489872 | 1.05739511 | 2.11773648 | 0.214581611 | 0.2524142244 |
| S. mystax | AMNH_188173 | 0.03922473 | 0.03600135 | 0.18469391 | 1.11973299 | 1.05514965 | 2.10158784 | 0.2606234645 | 0.2264232683 |
| S. mystax | AMNH_188177 | 0.0693735 | 0.06131289 | 0.19172642 | 1.14331347 | 1.05936652 | 1.94719523 | 0.3932765067 | 0.1018531273 |
| S. mystax | AMNH_188178 | 0.04486076 | 0.04167675 | 0.18895839 | 1.12580681 | 1.05872056 | 1.93388858 | 0.1874763787 | 0.1159032459 |
| S. imperator | FMNH_98035 | 0.03250676 | 0.03043884 | 0.15793409 | 1.11866975 | 1.05480596 | 2.24000748 | 0.1444650706 | 0.2952585792 |
| S. imperator | FMNH_98036 | 0.02248276 | 0.02094895 | 0.20258895 | 1.13024434 | 1.06076584 | 2.11511061 | 0.290536481 | 0.1745774121 |
| S. imperator | FMNH_121551 | 0.02670999 | 0.0255503 | 0.19245365 | 1.13287305 | 1.06052225 | 2.12716714 | 0.326739234 | 0.2095902606 |
| S. midas | FMNH_93516 | 0.02256071 | 0.02063783 | 0.20311824 | 1.12422849 | 1.05831849 | 2.16550041 | 0.2170084619 | 0.2583197171 |

**Table S4:** At the four studied epiphyses, for each variable (both topological indices and traditional parameters) and for the specimens representing each species (if there were data for at least 2 individuals) we computed the mean (x̄), the standard deviation (SD) and the coeffient of variation (CV).

**Proximal Humerus**

| | NodDen_mean | | | NodDen_median | | | TrabLen | | | TrabTort_mean | | | TrabTort_median | | | FD | | | DA | | | BV/TV | | |
|---|---|---|---|---|---|---|---|---|---|---|---|---|---|---|---|---|---|---|---|---|---|---|---|---|
| | x̄ | SD | CV | x̄ | SD | CV | x̄ | SD | CV | x̄ | SD | CV | x̄ | SD | CV | x̄ | SD | CV | x̄ | SD | CV | x̄ | SD | CV |
| *Leontocebus nigrifrons* | 0.015 | 0.002 | 0.107 | 0.014 | 0.002 | 0.131 | 0.206 | 0.010 | 0.049 | 1.126 | 0.009 | 0.008 | 1.060 | 0.003 | 0.002 | 2.122 | 0.074 | 0.035 | 0.279 | 0.039 | 0.141 | 0.130 | 0.043 | 0.332 |
| *Saguinus mystax* | 0.016 | 0.001 | 0.062 | 0.016 | 0.001 | 0.081 | 0.225 | 0.001 | 0.005 | 1.123 | 0.004 | 0.004 | 1.060 | 0.000 | 0.000 | 2.094 | 0.011 | 0.005 | 0.321 | 0.071 | 0.220 | 0.191 | 0.006 | 0.031 |
| *Saguinus imperator* | 0.009 | 0.000 | 0.024 | 0.009 | 0.000 | 0.015 | 0.211 | 0.012 | 0.055 | 1.113 | 0.002 | 0.002 | 1.060 | 0.001 | 0.001 | 2.236 | 0.086 | 0.039 | 0.305 | 0.011 | 0.037 | 0.183 | 0.009 | 0.051 |
| *Saguinus midas* | 0.009 | 0.002 | 0.230 | 0.009 | 0.002 | 0.227 | 0.230 | 0.011 | 0.046 | 1.116 | 0.000 | 0.000 | 1.058 | 0.001 | 0.001 | 2.102 | 0.075 | 0.036 | 0.282 | 0.007 | 0.024 | 0.173 | 0.027 | 0.158 |

**Distal Humerus**

| | NodDen_mean | | | NodDen_median | | | TrabLen | | | TrabTort_mean | | | TrabTort_median | | | FD | | | DA | | | BV/TV | | |
|---|---|---|---|---|---|---|---|---|---|---|---|---|---|---|---|---|---|---|---|---|---|---|---|---|
| | x̄ | SD | CV | x̄ | SD | CV | x̄ | SD | CV | x̄ | SD | CV | x̄ | SD | CV | x̄ | SD | CV | x̄ | SD | CV | x̄ | SD | CV |
| *Leontocebus nigrifrons* | 0.049 | 0.013 | 0.256 | 0.046 | 0.012 | 0.253 | 0.184 | 0.000 | 0.002 | 1.160 | 0.013 | 0.011 | 1.064 | 0.004 | 0.004 | 2.080 | 0.057 | 0.028 | 0.451 | 0.087 | 0.192 | 0.207 | 0.039 | 0.190 |
| *Saguinus mystax* | 0.078 | 0.009 | 0.120 | 0.074 | 0.009 | 0.123 | 0.205 | 0.004 | 0.018 | 1.175 | 0.013 | 0.011 | 1.071 | 0.004 | 0.004 | 1.780 | 0.073 | 0.041 | 0.498 | 0.020 | 0.041 | 0.171 | 0.017 | 0.099 |
| *Saguinus imperator* | 0.029 | 0.007 | 0.230 | 0.028 | 0.006 | 0.206 | 0.190 | 0.017 | 0.087 | 1.134 | 0.008 | 0.007 | 1.061 | 0.002 | 0.002 | 2.162 | 0.038 | 0.018 | 0.389 | 0.084 | 0.217 | 0.256 | 0.025 | 0.096 |
| *Saguinus midas* | 0.025 | 0.002 | 0.097 | 0.024 | 0.003 | 0.111 | 0.200 | 0.022 | 0.110 | 1.147 | 0.006 | 0.005 | 1.062 | 0.002 | 0.002 | 2.060 | 0.093 | 0.045 | 0.500 | 0.012 | 0.025 | 0.218 | 0.024 | 0.111 |

**Proximal Tibia**

| | NodDen_mean | | | NodDen_median | | | TrabLen | | | TrabTort_mean | | | TrabTort_median | | | FD | | | DA | | | BV/TV | | |
|---|---|---|---|---|---|---|---|---|---|---|---|---|---|---|---|---|---|---|---|---|---|---|---|---|
| | x̄ | SD | CV | x̄ | SD | CV | x̄ | SD | CV | x̄ | SD | CV | x̄ | SD | CV | x̄ | SD | CV | x̄ | SD | CV | x̄ | SD | CV |
| *Leontocebus nigrifrons* | 0.015 | 0.002 | 0.140 | 0.014 | 0.001 | 0.069 | 0.190 | 0.016 | 0.082 | 1.129 | 0.019 | 0.017 | 1.061 | 0.004 | 0.004 | 2.148 | 0.050 | 0.023 | 0.316 | 0.006 | 0.019 | 0.170 | 0.055 | 0.325 |
| *Saguinus mystax* | 0.016 | 0.002 | 0.123 | 0.015 | 0.002 | 0.106 | 0.222 | 0.006 | 0.029 | 1.120 | 0.009 | 0.008 | 1.061 | 0.001 | 0.001 | 2.117 | 0.037 | 0.018 | 0.359 | 0.107 | 0.297 | 0.218 | 0.021 | 0.097 |
| *Saguinus imperator* | 0.013 | 0.004 | 0.305 | 0.012 | 0.003 | 0.225 | 0.191 | 0.037 | 0.191 | 1.109 | 0.009 | 0.008 | 1.057 | 0.005 | 0.005 | 2.228 | 0.065 | 0.029 | 0.313 | 0.059 | 0.190 | 0.222 | 0.036 | 0.161 |

**Distal Tibia**

| | NodDen_mean | | | NodDen_median | | | TrabLen | | | TrabTort_mean | | | TrabTort_median | | | FD | | | DA | | | BV/TV | | |
|---|---|---|---|---|---|---|---|---|---|---|---|---|---|---|---|---|---|---|---|---|---|---|---|---|
| | x̄ | SD | CV | x̄ | SD | CV | x̄ | SD | CV | x̄ | SD | CV | x̄ | SD | CV | x̄ | SD | CV | x̄ | SD | CV | x̄ | SD | CV |
| *Leontocebus nigrifrons* | 0.039 | 0.004 | 0.102 | 0.036 | 0.004 | 0.113 | 0.183 | 0.004 | 0.021 | 1.130 | 0.006 | 0.005 | 1.059 | 0.002 | 0.002 | 2.078 | 0.038 | 0.018 | 0.219 | 0.023 | 0.107 | 0.190 | 0.066 | 0.344 |
| *Saguinus mystax* | 0.051 | 0.016 | 0.313 | 0.046 | 0.013 | 0.287 | 0.188 | 0.004 | 0.019 | 1.130 | 0.012 | 0.011 | 1.058 | 0.002 | 0.002 | 1.994 | 0.093 | 0.047 | 0.280 | 0.104 | 0.372 | 0.148 | 0.068 | 0.461 |
| *Saguinus imperator* | 0.027 | 0.005 | 0.185 | 0.026 | 0.005 | 0.185 | 0.184 | 0.023 | 0.127 | 1.127 | 0.008 | 0.007 | 1.059 | 0.003 | 0.003 | 2.161 | 0.069 | 0.032 | 0.254 | 0.096 | 0.380 | 0.226 | 0.062 | 0.274 |

**Note S2:** We oriented the humeri following Appendix Fig. 2 of Ruff (2002). In brief, the bone is first placed with the posterior surface (highlighted in orange in Fig. A), pointing down. To do this, we observed the bone in distal view (Fig. A) and we rotated it in the transverse plane (defined by the x and y axis highlighted in green in Fig. A) until the mid-points of the lateral and medial epicondyles were lying along the x axis. In this way, the tibia is oriented in the transverse plane. To orient the tibiae in the coronal plane (defined by the x and z axis highighted in green in Fig. B, showing the bone observed from a posterior view), we made the mid-points of the mediolateral (ML) distance on the shaft in correspondence of the surgical neck (shown proximally in Fig. B, with two intersecting blue arrows) and the point of intersection of the trochlea and the capitulum (approximately corresponding to the point where the y axis intersects the bone on the anterior surface in Fig. A, highlighted with a black ellipse) to intersect the z-axis. Similarly, to orient the humeri in the sagittal plane (which is perpendicular to the coronal. It is not shown here but, similarly to the coronal plane, it shows the bone in its longest size, i.e. proximodistal) we made the mid-point of the anteroposterior distance on the shaft in correspondence of the surgical neck and the mid-point and the AP distance in correspondence of the lateral lip of the trochlea (approximately corresponding to the purple ellipse in Fig. A) to insersect the z axis.

**A. Distal view**

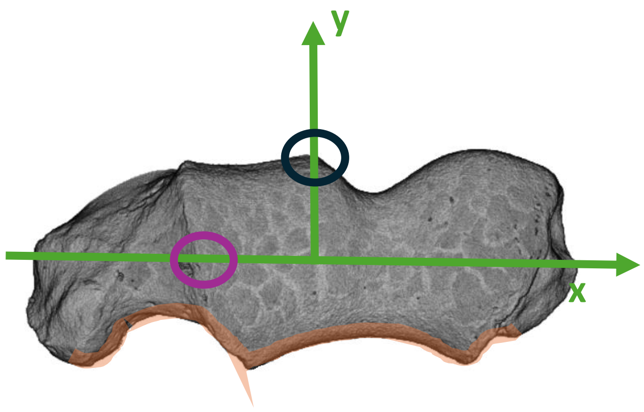


**B. Posterior view**

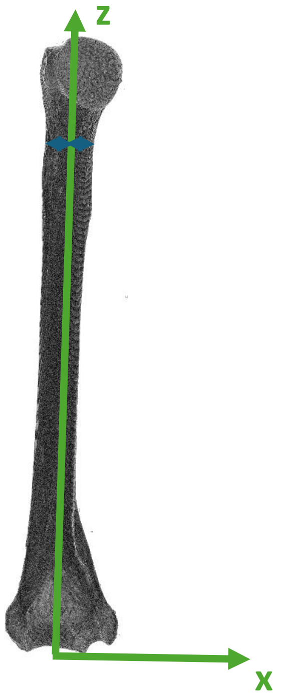

**Note S3:** We oriented the tibiae following Appendix Fig. 2 of Ruff (2002). In brief, the bone is first placed with the posterior surface (highlighted in orange in Fig. A),  pointing down. To do this, we observed the bone in proximal view (Fig. A) and we rotated it in the transverse plane (defined by the x and y axis highlighted in green in Fig. A) until the mid-points of the anteroposterior (AP) lengths of the lateral and medial condyles were at the same distance from the x axis. In Figure A, the medial and lateral condyles are highlighted in light blue, the AP lengths of the lateral and medial condyles are represented by black dashed lines, their mid-points are represented by the intersection between the black dashed lines and the horizontal yellow dashed line. In other words, orienting the tibia following these criteria, corresponds to make the AP lengths parallel to the y axis and the yellow dashed line parallel to the x axis. In this way, the tibia is oriented in the transverse plane. To orient the tibiae in the coronal plane (defined by the x and z axis highighted in green in Fig. B, showing the bone observed from a posterior view) we made the mid-points of the proximal and distal tibial plateaus (represented by the interesection of the blue arrows, both proximally and distally) to intersect the z axis (Fig. B). Similarly, to orient the tibiae in the sagittal plane (which is perpendicular to the coronal. It is not shown here but, similarly to the coronal plane, it shows the bone in its longest size, i.e. proximodistal) we made the mid-point of the proximal tibial plateau (approximately corresponding to the intersection between the y axis and the yellow dashed line in Fig. A) and the mid-point of the tibio-talar surface (shown with the interserction of the two yellow arrows in Fig. C, while the surface is highlighted in blue) to intersect the z-axis.

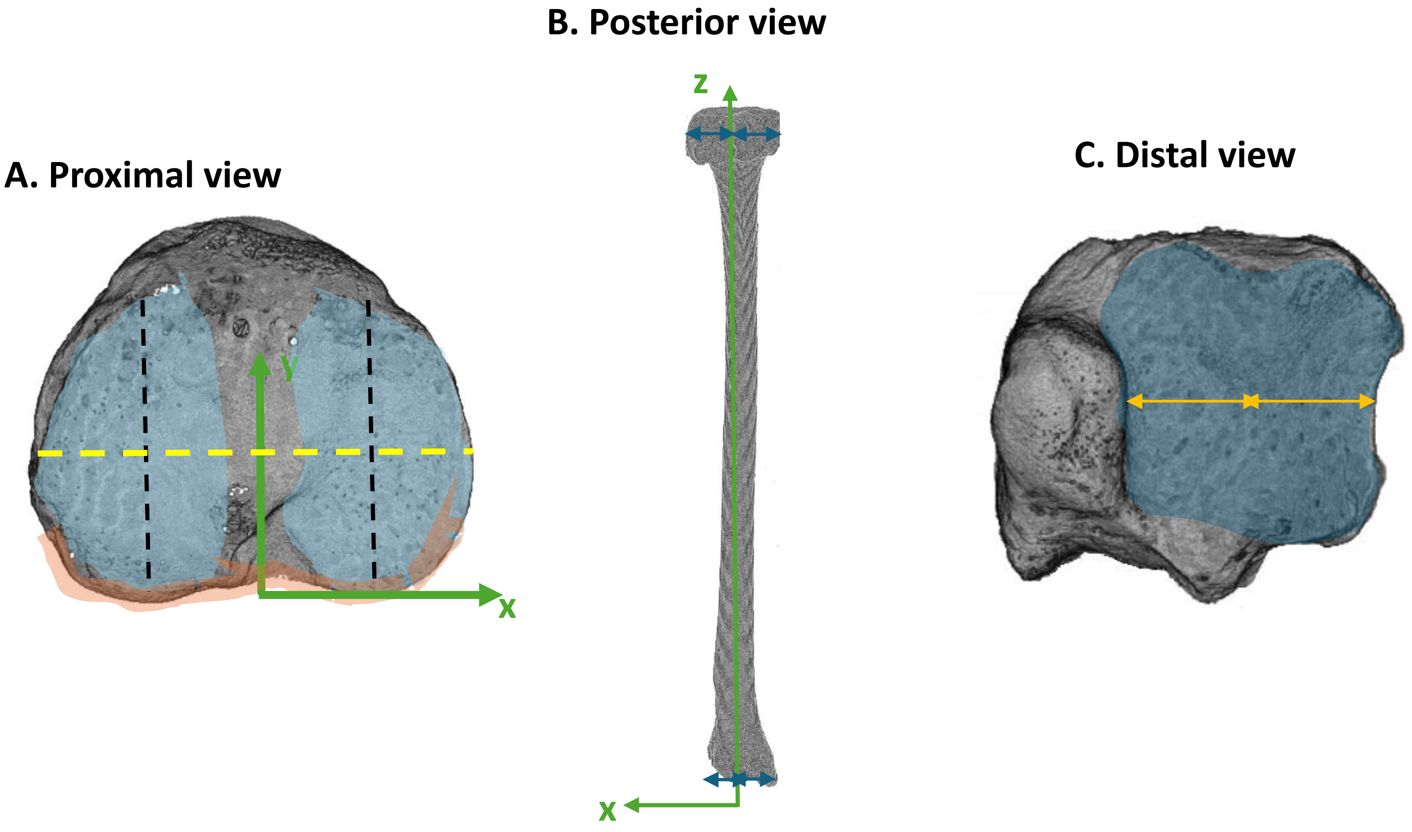

**Note S4:** The variation of the trabecular variables studied in this work is shown for the sampled specimens divided by sex, through PCAs performed at each humeral/tibial epiphysis. PC1-PC2 and PC2-PC3 biplots are shown.

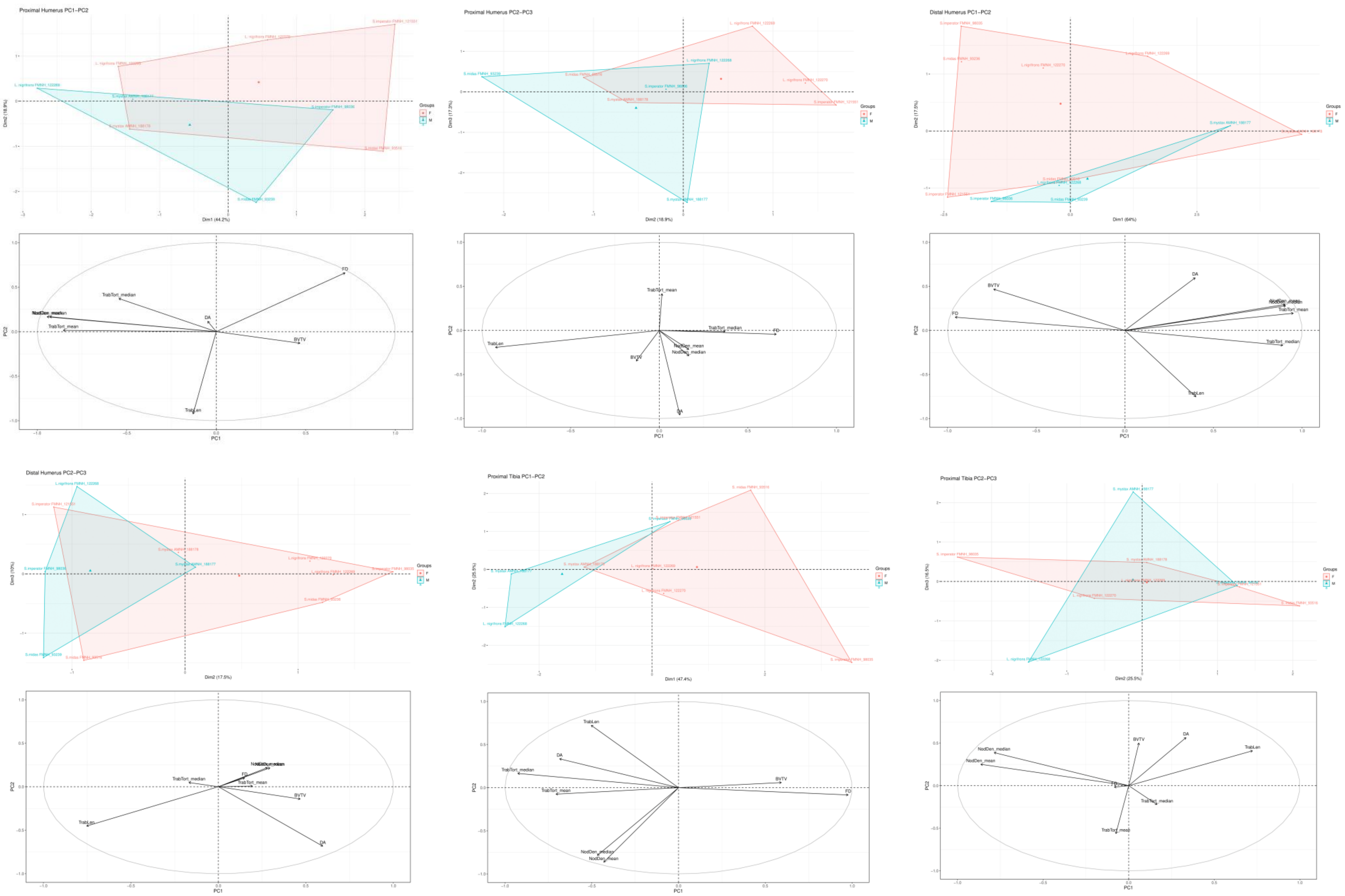

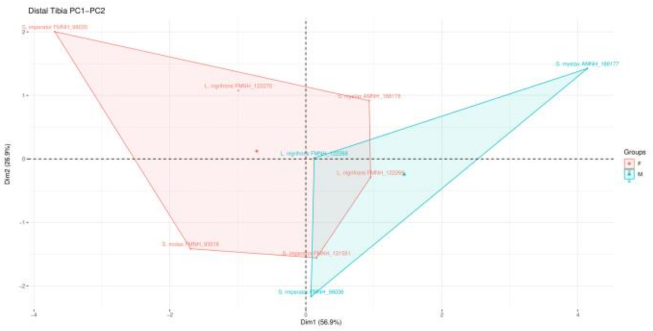


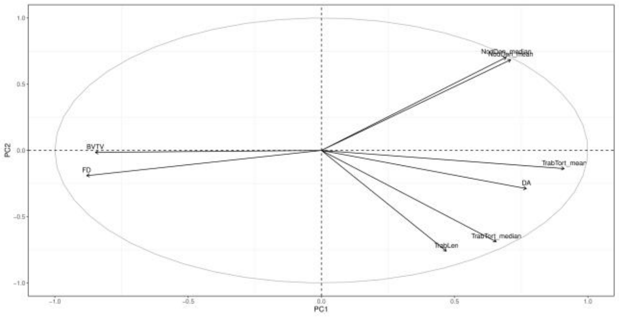


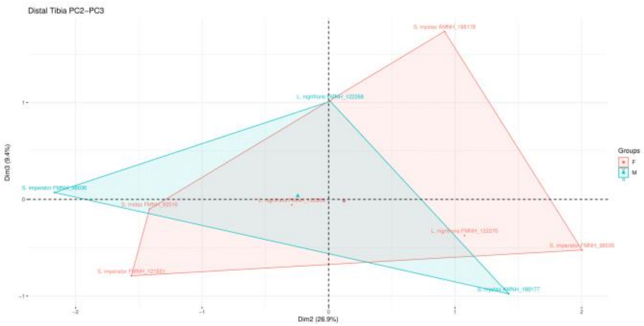


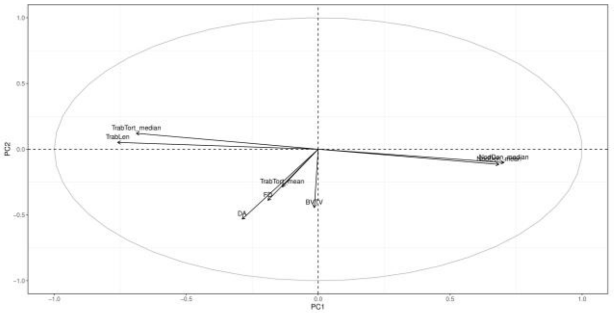


- A trend of relative increase in FD for female individuals can be detected in the proximal humerus (as one can evince from female individuals tending to occupy the upper right and right regions of the PC1-PC2 and PC2-PC3 morphospaces, respectively, together with the respective variable loading plots), distal humerus (as one can evince from female individuals tending to occupy the right region of the PC1-PC2 morphospace, together with the respective variable loading plot), proximal tibia (as one can evince from female individuals tending to occupy the right region of the PC1-PC2 morphospace, together with the respective variable loading plot) and distal tibia (as one can evince from female individuals tending to occupy the left region of the PC1-PC2 morphospace, together with the respective variable loading plot)

- A trend of relative increase in TrabLen for male individuals can be detected in the proximal humerus (as one can evince from male individuals tending to occupy the left region of the PC2-PC3 morphospace, together with the respective variable loading plot), distal humerus (as one can evince from male individuals tending to occupy the bottom right region of the PC1-PC2 morphospace, together with the respective variable loading plot)

- A trend of relative increase in BV/TV for female individuals can be detected in the proximal tibia (as one can evince from female individuals tending to occupy the right region of the PC1-PC2 morphospace, together with the respective variable loading plot) and distal tibia (as one can evince from female individuals tending to occupy the left region of the PC1-PC2 morphospace, together with the respective variable loading plot)

- A trend of relative increase in DA for male individuals can be detected in the proximal tibia (as one can evince from male individuals tending to occupy the left region of the PC1-PC2 morphospace, together with the respective variable loading plot) and distal tibia (as one can evince from male individuals tending to occupy the right region of the PC1-PC2 morphospace, together with the respective variable loading plot)

- A trend of relative increase in TrabTort for male individuals can be detected in the proximal tibia (as one can evince from male individuals tending to occupy the left region of the PC1-PC2 morphospace, together with the respective variable loading plot) and partially, i.e. mainly through TrabTortMean, in the distal tibia (as one can evince from male individuals tending to occupy the right region of the PC1-PC2 morphospace, together with the respective variable loading plot)

**Note S5:** The variation of the trabecular variables studied in this work is shown for the sampled specimens divided by captivity, through PCAs performed at each humeral/tibial epiphysis. PC1-PC2 and PC2-PC3 biplots are shown.

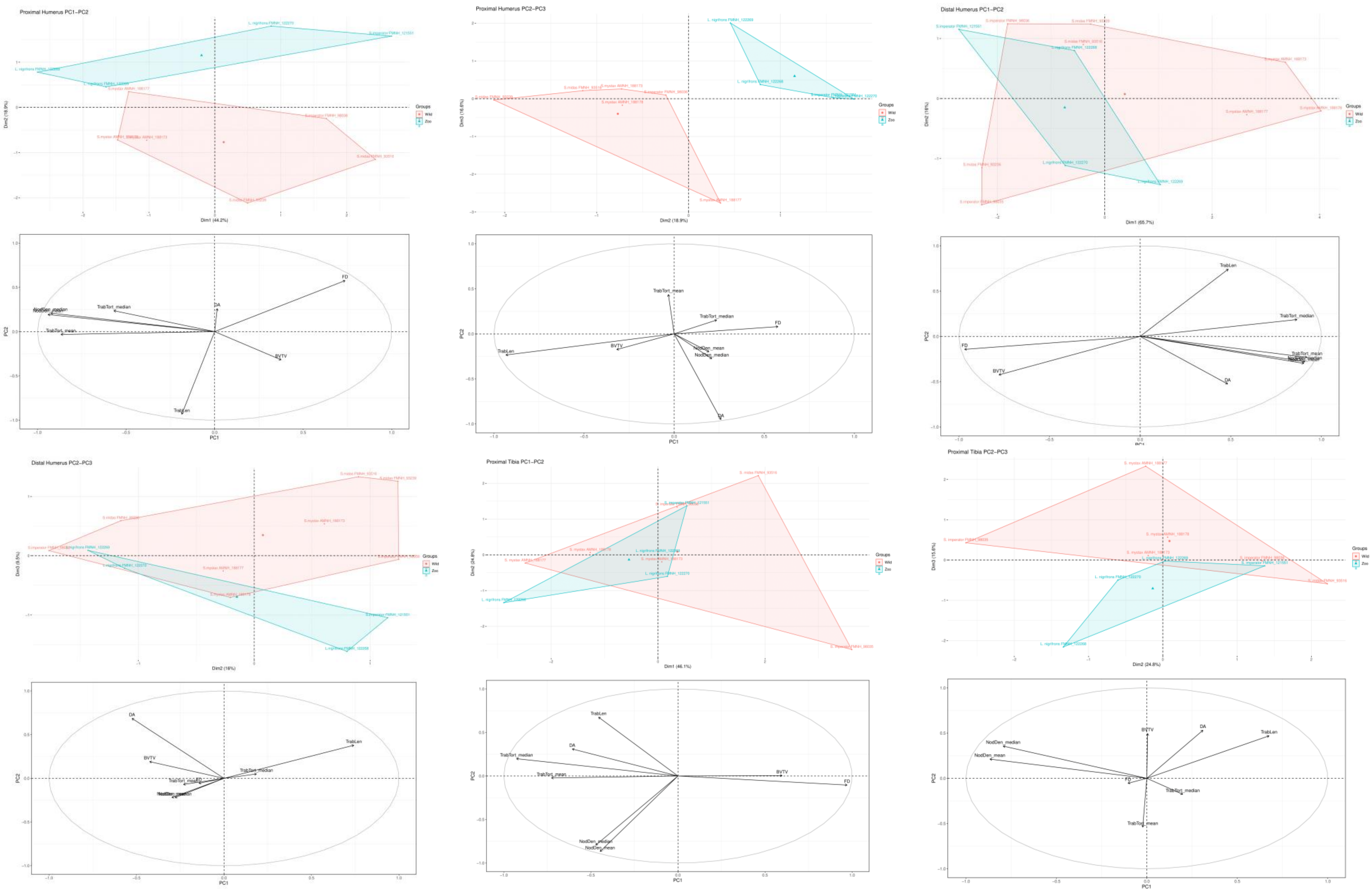

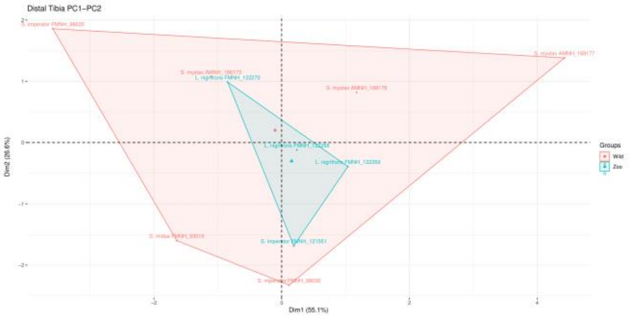


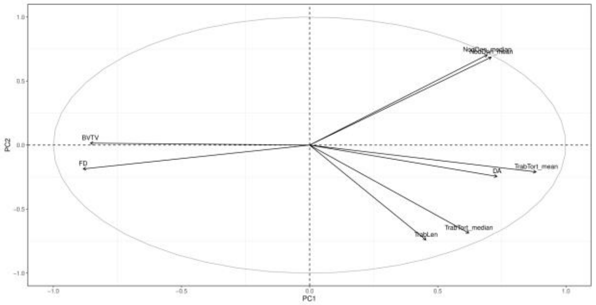


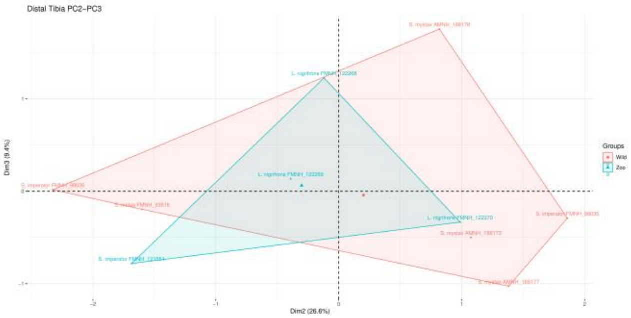


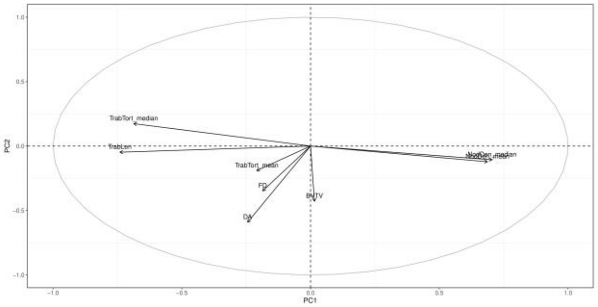


- A trend of decrease in TrabLen for captive individuals can be detected in the proximal humerus (as one can evince from captive individuals tending to occupy the upper region and the upper right region of the PC1-PC2 and PC2-PC3 morphospaces, respectively, together with the respective variable loading plot)

- A trend of relative increase in TrabTort for captive individuals can be detected in the proximal tibia (as one can evince from captive individuals that are driven by TrabTort toward the bottom region of the PC1-PC2 morphospace, although with minor contribution)

- A trend of relative decrease in BV/TV for captive individuals can be detected in the proximal humerus (as one can evince from captive individuals tending to occupy the upper right region in the PC2-PC3 morphospace, together with the respective variable loading plot) and the proximal tibia (as one can evince from captive individuals tending to occupy the bottom region in the PC1-PC2 morphospace, together with the respective variable loading plot)

- A trend of relative increase in FD for captive individuals can be detected in the proximal humerus (as one can evince from captive individuals tending to occupy the upper right region in the PC2-PC3 morphospace, together with the respective variable loading plot)

**Note S6-S9:** Heat maps for NodDen distribution  are shown through 2D sections going proximodistally in the epiphyses at various levels of the total epiphyseal length (expressed in %).

**Note S6: Proximal Humerus NodDen maps (long leapers in blue, short leapers in light red)**

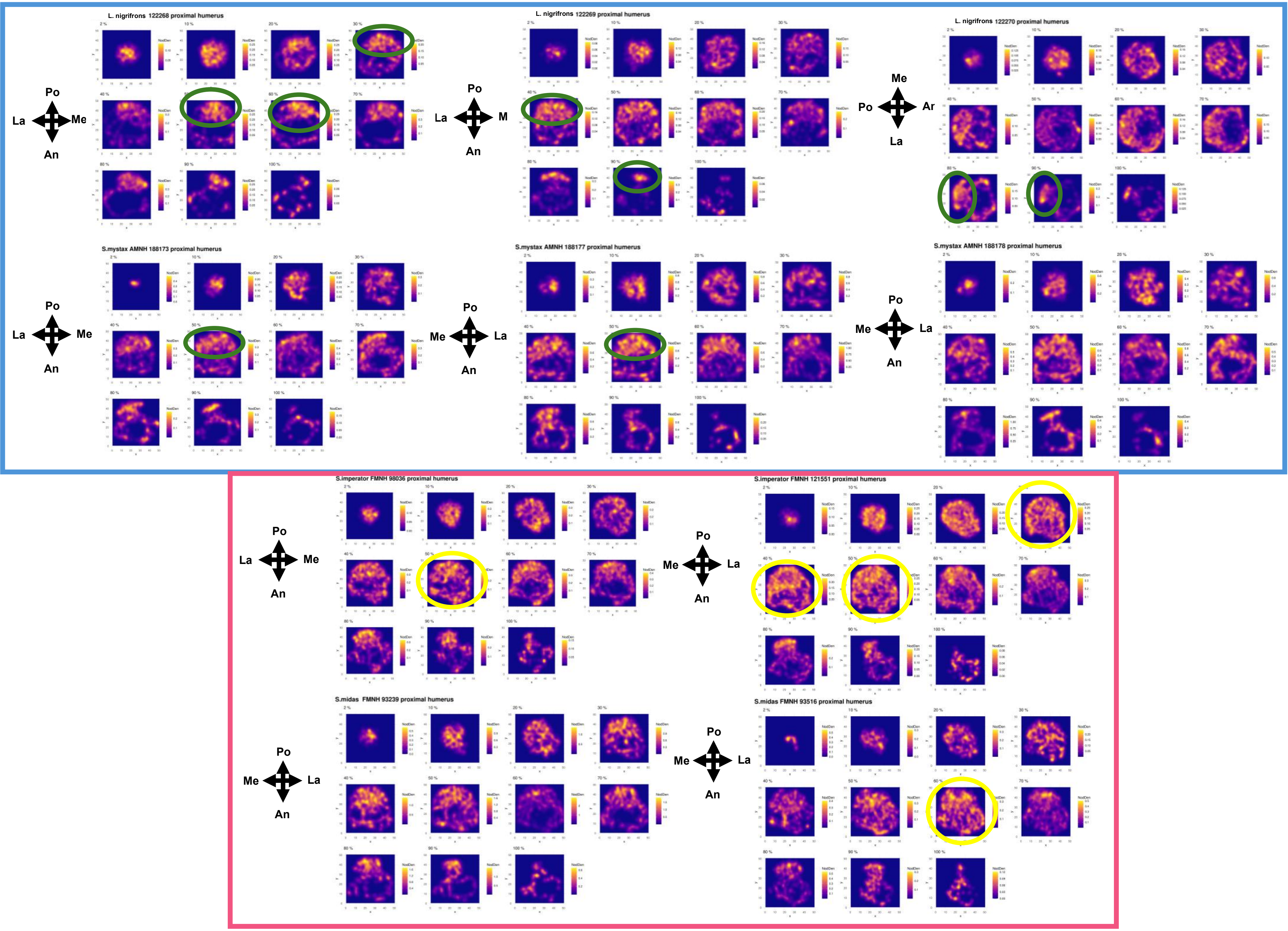

#### Note S7: Distal Humerus NodDen maps

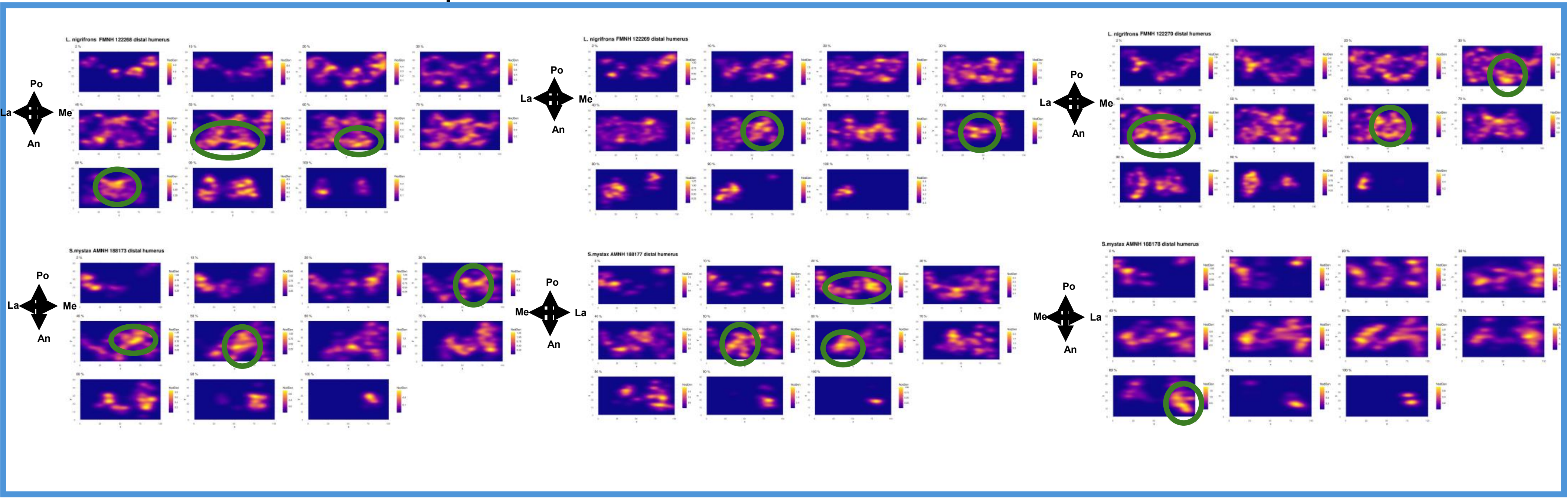


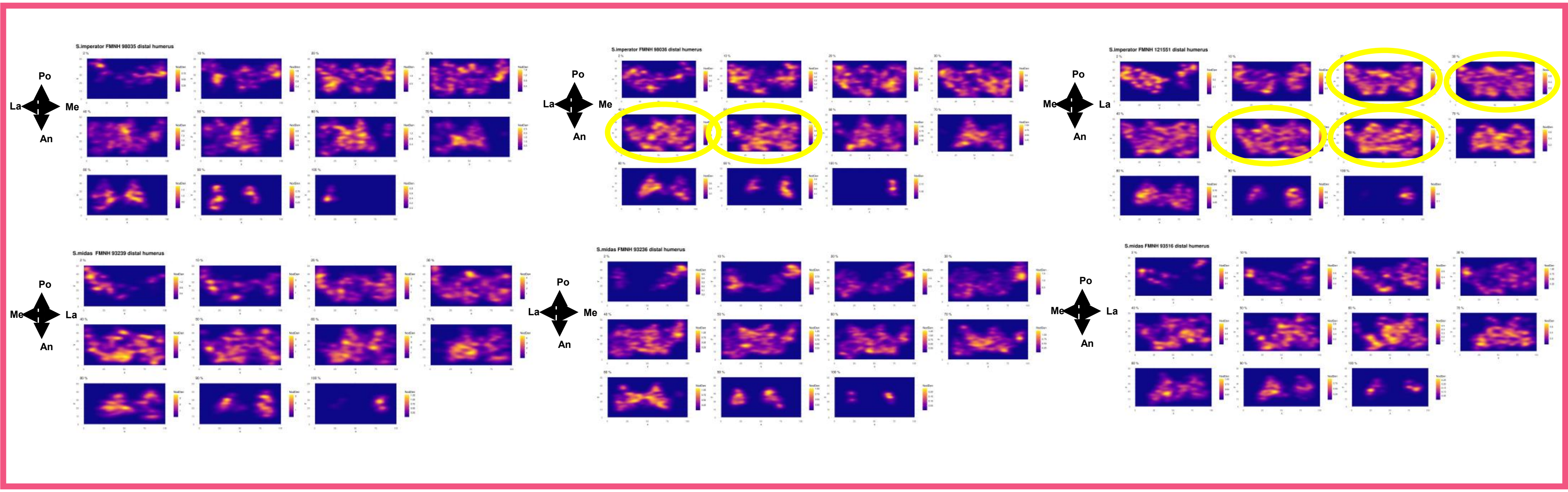

### Note S8: Proximal Tibia NodDen maps

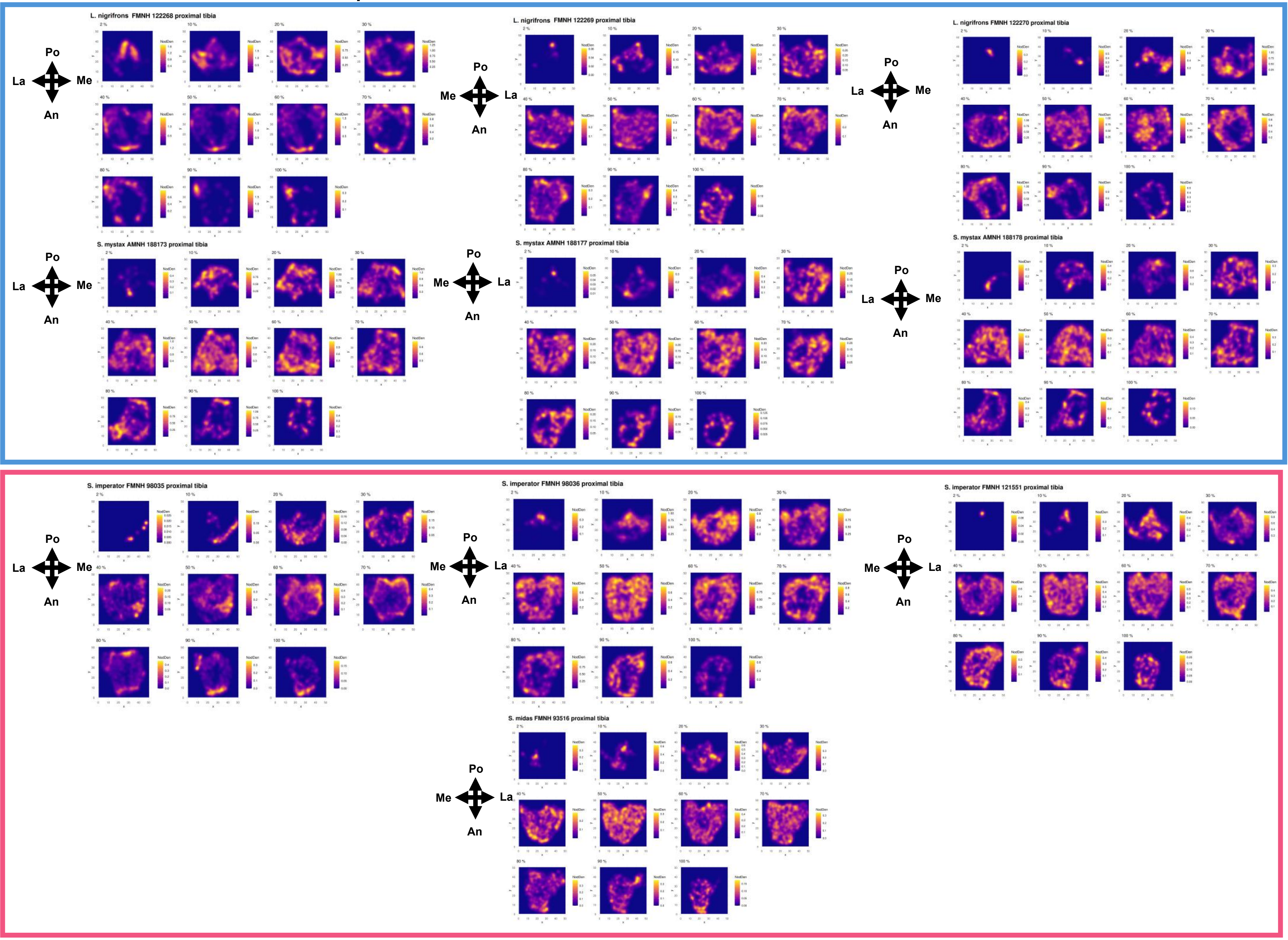

### Note S9: Distal Tibia NodDen maps

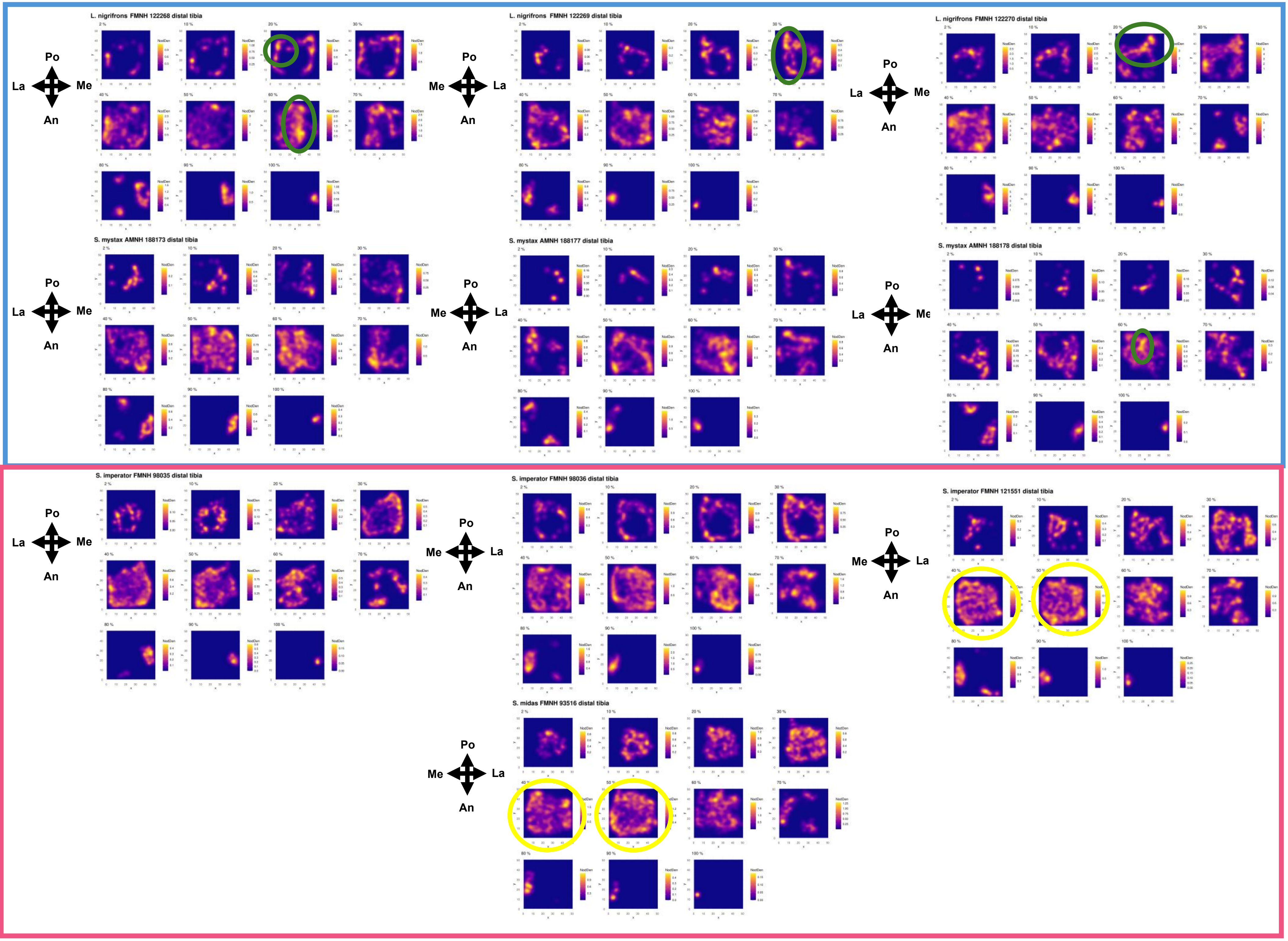
